# IFNγ insufficiency during mouse intravaginal *Chlamydia trachomatis* infection exacerbates alternative activation in macrophages with compromised CD40 functions

**DOI:** 10.1101/2022.07.05.497583

**Authors:** Naveen Challagundla, Dhruvi Shah, Sarat K Dalai, Reena Agrawal-Rajput

## Abstract

*Chlamydia trachomatis (C*.*tr)*, an obligate intracellular pathogen, causes asymptomatic genital infections in women and is also a leading cause of preventable blindness. Limited mouse model of chronic *C*.*tr* infection are available to study the host immune response. We have developed *in vivo* mice models of acute and chronic infections for *C. trachomatis* to explore the significance of macrophage-directed response in mediating immune activation/suppression. During chronic and repeated infections, IFNγ secretion from T cells is abated while TGFβ and IL-10 secretion is enhanced. An increase in exhaustion (*PD1, CTLA4*) and anergic (*Klrg3, Tim3*) T cell markers is also observed during chronic infection. It was observed that alternatively-activated macrophages with low CD40 expression promote Th2 and Treg differentiation and lead to sustained *C. trachomatis* genital infection. Macrophages infected with *C. trachomatis* or treated with supernatant of infected epithelial cells drive them to alternately-activated phenotype. *C. trachomatis* infection prevents increase in CD40 expression. Low IFNγ, as observed in chronic infection leads to incomplete clearance of bacteria and poor immune activation. *C. trachomatis* decapacitates IFNγ responsiveness in macrophages via hampering *IFNγRI* and *IFNγRII* expression which can be correlated with poor expression of MHC-II, CD40, iNOS and NO release even following IFNγ supplementation. Alternatively-activated macrophages during *C. trachomatis* infection express low CD40 rendering immunosuppressive, Th2 and Treg differentiation which could not be reverted even after IFNγ supplementation. The alternative macrophages also harbour high bacterial load and are poor responders to IFNγ, thus promoting immunosupression. Thus, *C. trachomatis* modulate the innate immune cells attenuating the anti-chlamydial functions of T cells in a manner that involves decreased CD40 expression on macrophages.

## Introduction

Female genital tract maintains a tightly regulated immune response that balances reproductive tolerance and protection against genital pathogenicity. Inflammatory responses are necessary to effectively eliminate pathogens, but uncontrolled inflammation has been reported to enhance the risk of disease acquisition and progression to pathological conditions. *C. trachomatis* (*C*.*tr*) binds to host surface receptors inducing endocytic uptake into membrane bound vesicles, where it escapes the fusion to lysosome which allows its replication/persistence inside the vesicle by utilizing the host metabolites [1]. *C*.*tr* primarily infects epithelial and endothelial cells, however, *C*.*tr* released from epithelial cells by extrusion or lysis can be phagocytosed by macrophages. Sustained inflammation during sexually transmitted *C*.*tr* infection causes pelvic inflammatory disease and long-term consequences like infertility, ectopic pregnancy, fallopian tube scarring and is reported to facilitate the transmission of human immunodeficiency virus [2]. *C*.*tr* can undergo spontaneous clearance in the host or may persist for months and progress as latent infection. The clearance or persistence of infection is presumed to be an outcome of varied immune response. The mechanism regarding the failure in the clearance of *C*.*tr* is poorly explained, but it might be the consequence of immune evasion strategies of the organism which may result in the failure to mount an appropriate immune response [3].

In addition to other co-stimulatory signals, it is the interaction of CD40-CD40L and release of IL-12 from the APCs, that activate CD4^+^T cells to differentiate into interferon-γ (IFNγ) producing Th1 cells. The CD8^+^ T cells are primed against *C*.*tr* infection via MHC-I and the IFNγ release from the Th1 cells also aid in this activation contributing to clearance of infection. Further, IFNγ activates downstream signals like iNOS, SOCS, STAT1, indolamine 2,3-dioxygenase (IDO) even in the innate immune cells contributing to clearance of pathogen [4, 5]. IFNγ downregulates IDO to cause tryptophan depletion which inhibits the growth of *C. trachomatis*. Some of the genital serotype of *C. trachomatis* may express *trpB/A* that converts indole to tryptophan mediating IFNγ-IDO resistance resulting in resistance to host defence [6]. Even though CD40-CD40L interaction is necessary for IFNγ production, studies have reported no significant change in CD40 expression in infected macrophages or dendritic cells in the genitals [7, 8]. However, CD40 is reported to be essential for conferring the generation of effective CD8^+^ T memory cells and protective immunity [9, 10]. Even though CD4^+^ T cells help in controlling chlamydial infection, little is known about the mechanism of persistent infection and *C*.*tr* induced immunosuppression.

Macrophages demonstrate varied plasticity, where they may exist as non-activated (M_0_), classically-activated (M_1_) or alternatively-activated (M_2_) macrophages. Different macrophage phenotypes led to differential responsiveness to infection, so the functional outcomes are diverse. Naive macrophages, following uptake of pathogens, get activated to M_1_ macrophage which release inflammatory cytokines and recruit T cells to activate them for clearance of bacteria. However, macrophages may also secrete anti-inflammatory cytokines during alternative activation. The classical/alternatively activated macrophage at the site of infection dictates the immune outcome. It has been previously reported that anti-inflammatory immune cells like M_2_ and Th2 exist in abundance at the site of *Chlamydia* infection in women [11]. The presented article aims at unravelling the mechanism of immune activation/suppression of macrophages during *C*.*tr* infection that may dictate the outcome of immune response via influencing the T cell response.

## Results

### Repeated vaginal infection of C.tr leads to persistent infection compromising immune response

Balb/c mice were infected repeatedly every 3 days to mimic: [chronic (C), chronic infection resolved (CR) or secondary infection (SI)]. The other groups included infection only once to mimic: [acute (A) or acute infection resolved (AR)] group as mentioned in Fig.1A. *C*.*tr* burden was analysed every two days for first 30 days and every 5 days for next 30 days. Bacterial shedding was analysed in cervical swab, the bacterial burden in cervical lavage and cervix tissue were taken as an indicator of infection status at the cervix. The chronic infection group (C) showed the presence of *C*.*tr* as observed by bacterial shedding and bacterial burden in cervical lavage and cervical tissue. Chronic resolved (CR) group of mice did not show *C*.*tr* shedding by day 45, however, bacterial burden was seen in the cervical lavage and cervical tissue (Fig.1B-D, supplementary figure (SF). 1). In secondary infection group (SI), infection was re-established as indicated by the presence of *C*.*tr* in cervical lavage, cervical tissue and bacterial shedding. The acute infection demonstrates high frequency of *C*.*tr* as observed by its presence in the swab, lavage and tissue. The acute resolved (AR) infection mice showed *C*.*tr* persistence as observed by presence of bacteria in swab, lavage and cervical tissue. These results indicate that chronic infection showed poor clearance of C.tr indicating poor mounting of recall response (Fig. 1B-D). Repeated infection groups (C, CR and SI) not only showed poor bacterial clearance, but also demonstrated fluid accumulation, hydrosalpinx as an indicator of inflammation (Fig. 1E). Infected mice show splenomegaly and increase in the size of inguinal lymph nodes compared to naïve mice (SF.2). Repeated infection groups (C, CR and SI) show tissue damage and aberrant growth of epithelial cells (Fig.1F). Acute infection groups (A and AR) demonstrated higher neutrophil recruitment. Chronic infection group (C) demonstrated higher neutrophil recruitment compared to lymphocyte which was diminished by day 60. Secondary infection showed poor neutrophil and lymphocyte recruitment and the ratio of neutrophil to lymphocyte to low. However, SI also demonstrated neutrophil infiltration confirming the establishment of infection. Despite high lymphocyte infiltration, the poor clearance of bacteria may be attributed to anti-inflammatory/ immunosuppressive T cell subset (Th2 and Treg; which is explored later in the manuscript) (Fig. 1G). Serum IFNγ levels monitored during the infection had high IFNγ levels early in infection, which rapidly decreased correlating with bacterial clearance. However, on repeated infections and secondary infection, IFNγ levels were not increased in serum and correlates with poor bacterial clearance indicating poor immune elicitation in chronic (C and CR) and secondary infections (SI) (Fig.1H). Acute infection showed increased serum IgG levels which decreased in AR group. Chronic resolved (CR) and secondary infection (SI) showed decreased serum IgG levels compared to acute infection (ARvsCR). IgA levels were increased in Acute infection which was later decreased (AR). Chronic and Secondary infections show IgA secretions however, they were lower compared to acute (A and AR) infections (Fig.1I and J). Tissue inflammatory status of the infection was analysed in the vaginal lavage of infected mice. The VL of acute infection groups (A and AR) had high levels of TNFα and IFNγ. However, the VL from chronic infected mice demonstrate the release of TNFα, but it accompanied increased release of IL-10. Interestingly, these group of mice failed to secrete the IFNγ in lavage which may be the justification for poor bacterial clearance. The resolving phase mice show increased release of immunosuppressive cytokine IL-10 and decrease release of TNFα. Secondary infection failed to release high IFNγ which is necessary for the bacterial clearance (SI). Similarly, enhanced release of IL-10 was observed that may be contributing to immunosuppression mediated infection establishment (Fig.1K).

**Figure 1:**
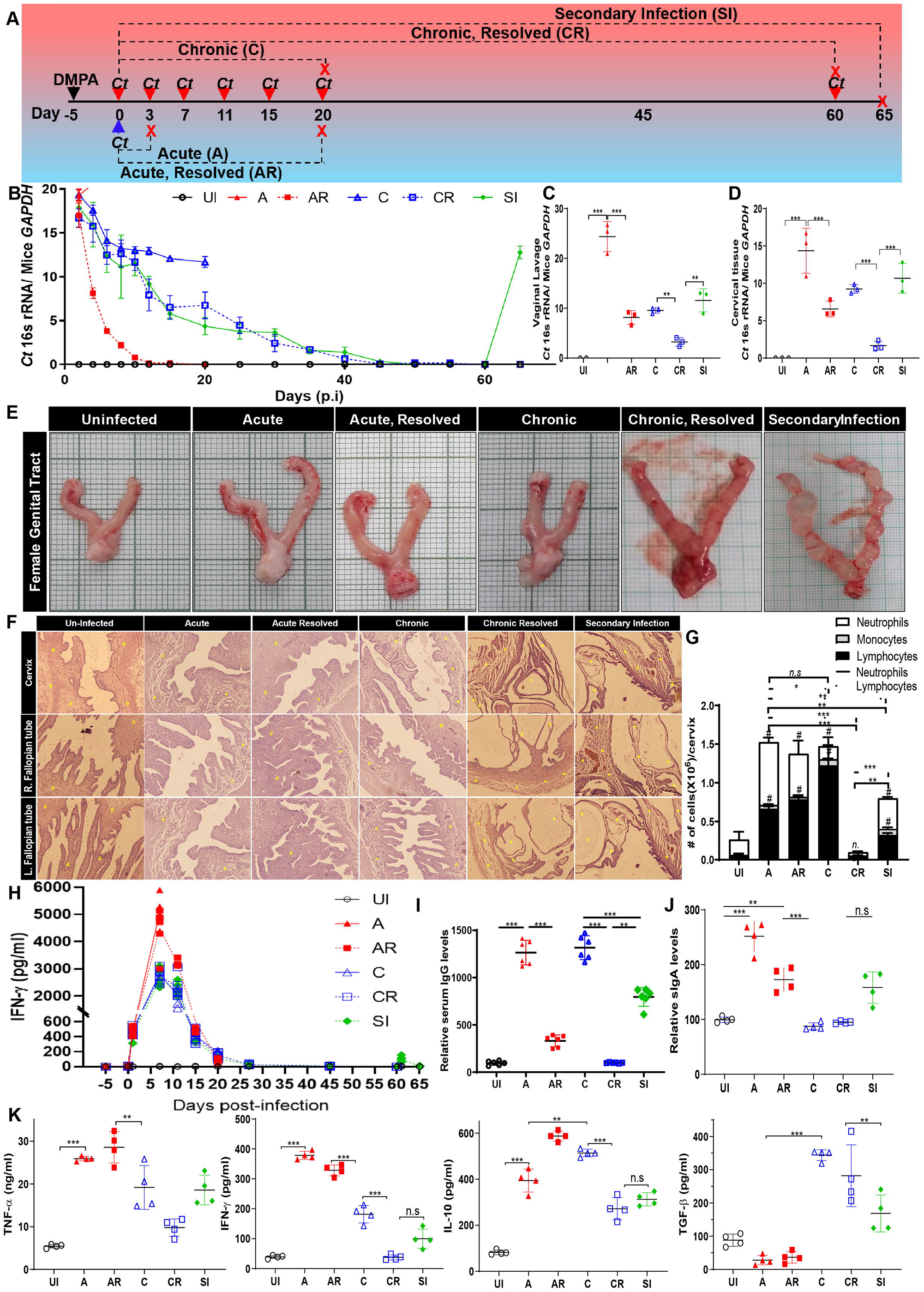
Chronic intravaginal *Chlamydia trachomatis* infection induces immune suppression and contribute to disease severity. Schematic representation of *C*.*tr* intra vaginal infection (A). *C*.*tr* burden were observed from cervical swab at indicated time points (B). *C*.*tr* burden was observed in cervical lavage and represented as ratio of DNA of *C*.*tr*:*mus musculus* (C). Total DNA was isolated from equal amount of tissues from all groups of mice and bacterial burden was analyzed as mentioned in A(D). Anatomical appearance of uninfected and infected female genital tract (E). H&E staining of the genital tract section of the infected and control animals (F). Vaginal lavage was collected from infected and control mice and analyzed for differential WBC count in VetScan and plotted as absolute number of cells per mice (G). Retro-orbitally blood was collected from control and infected mice. Serum separated from blood was used to measure IFNγ levels by ELISA (H). *C*.*tr*-specific IgG levels levels were measured form serum of infected and control mice by using direct cell-based ELISA and plotted as relative IgG levels (I). IgA levels were measured in the vaginal lavage by ELISA (J). Cytokines: TNFα, IFNγ, IL-10 and TGFβ were measured from the vaginal lavage by using ELISA (K). (n=6; ***p<0.0001; **p<0.001)

In order to establish the involvement of macrophages for the secreted IL-10, we analysed the macrophage population for M_1_/M_2_ markers by analysing CD68 and CD206 expression on CD11b macrophages respectively. The acute infection group demonstrated that the recruited macrophages were expressing high levels of CD68 while the other groups had significantly lower levels compared to acute infection group. The mice in the chronic infection groups (C and CR) and secondary re-infection (SI) show high levels CD206 (M_2_ marker) while the acute infection group were low in CD206 expression on macrophages (Fig.2A and B). The phenotypic markers corelates with iNOS expression (Fig.2D) and arginase activity (Fig.2E) in the cervical tissue indicating higher M_1_ macrophage recruitment during acute (A) infection while chronic (C and CR) and secondary infection group of mice (SI) show higher M_2_ macrophages. Since M_1_ and M_2_ macrophages expresses varied levels of CD40 and this differential CD40 activation dictates macrophage secretory output of IL-10 and IL-12, CD40 levels were analysed from cervical tissues of infected and uninfected mice by IHC. CD40 was poorly expressed in chronic and secondary infections on recruited CD11b^+^ macrophages (Fig. 2C and D). T cells recruited to the cite of infection plays a major role in clearance of infection by their ability to secrete IFNγ. Acute infection show higher CD4^+^ and CD8^+^ T cell recruitment compared to chronic and secondary infections (Fig.2 F and G). Chronic infection also shows higher CD4^+^ T cell recruitment compared to control but the resolving phase of infection demonstrated lower CD4^+^ T cell recruitment. Similarly, the acute infection group showed higher CD8^+^ T cell recruitment compared to chronic (C and CR) and secondary infection (SI) groups (Fig.2G). Further, T cell activation during the infection was analysed from the cervical tissue draining inguinal lymph nodes. On acute infection, mice show higher IFNγ producing CD4^+^ T and CD8^+^ T cells, which were significantly decreased during chronic infection (C and CR) and secondary re-infection (SI) compared to acute infection groups corelating with poor CD40 expression and anti-inflammatory macrophages at site of infection (Fig.2H). The T cells from acute infection (A) also show increased IFNγ and TNFα secretion which was decreased in resolving group of mice (AR). T cells from chronic infections and secondary infection failed to secrete both IFNγ and TNFα. Secondary infection group was able to elicit T cells to secret IFNγ, but it was very low comparatively (Fig. 2I). These T cells were analysed for co-stimulatory (*CD40L*) or inhibitory (*PD1*) and *T-bet* mRNA expression to co-relate and reason their activation state. T cells from chronic (C and CR) and secondary infection (SI) group of mice show poor CD40L and T-bet (Th1 transcription factor) mRNA levels. *C*.*tr* chronic infection increased PD1 expression on inguinal lymph node T cells (Fig.2J). To analyse the systemic immune response, splenic T cells activation state was analysed. Acute *C*.*tr* infected mice show high IFNγ^+^ CD4^+^ T cells. The chronic infection groups show decrease in IFNγ producing CD4^+^ T cells with simultaneous increase in TGFβ producing CD4^+^ T cells. Secondary infection also elicited more TGFβ producing CD4^+^ T cells compared to IFNγ producing CD4^+^ T cells (Fig.2K). This decrease in activation of IFNγ producing T cells leads to decreased IFNγ mediated bacterial clearance. The improper T cell activation could be due to skewed macrophage activation by *C. trachomatis*.

**Figure 2:**
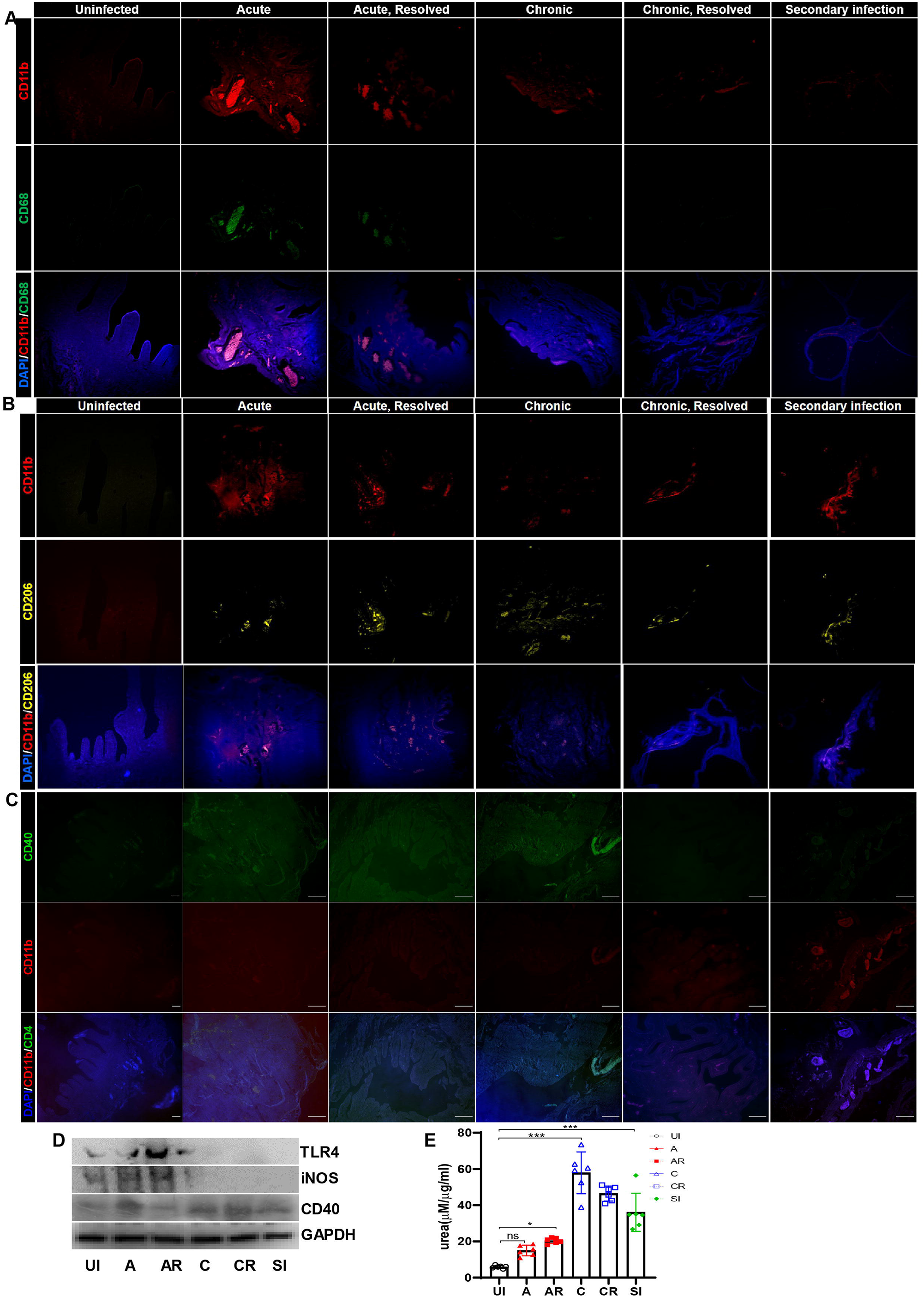
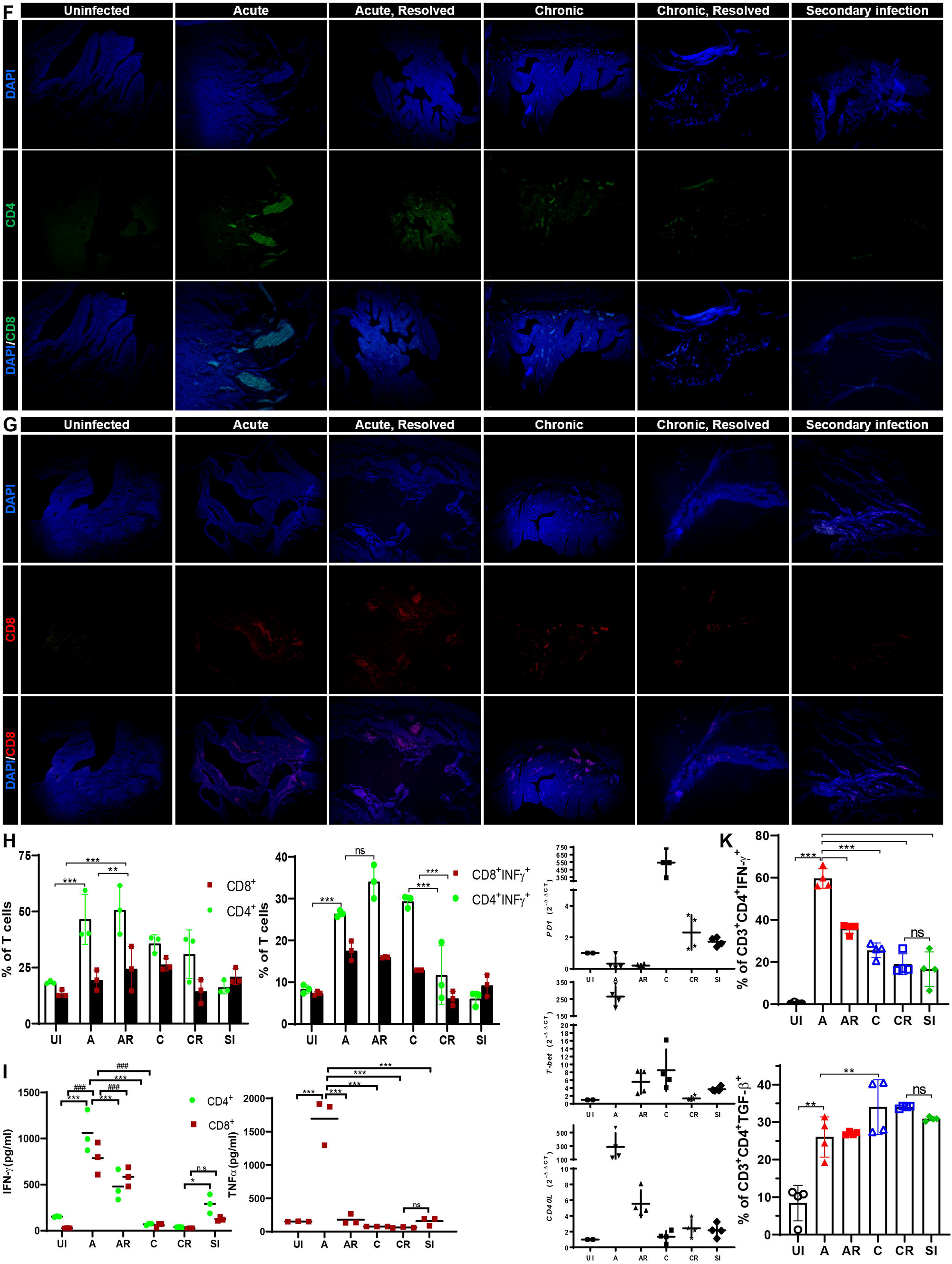
Chronic *C. trachomatis* infection suppresses IFNγ producing T cells. Section of the genital tract of infected and control mice were stained for CD40 using anti-CD40 which is identified using anti-rabbit IgG labelled with FITC. Simultaneously anti-CD11b labelled with APC was used to observe macrophages in background of nuclear stain, DAPI at 200X magnification (A). Immunohistochemistry was done for CD11b, CD68 using anti-CD11b labelled with APC and CD68 labelled with FITC to observed M_1_ macrophages (B). Immunohistochemistry was done for CD11b, CD206 using anti-CD11b labelled with APC and CD206 labelled with PE to observed M_2_ macrophages (C). CD40 and iNOS expression levels were observed using western blot from the RIPA lysed for tissue of infected and control mice (D). Simultaneously tissue was lysed in PBS used to measure arginase activity by measuring conversion of arginine to urea per ug of protein per ml (E). Immunohistochemistry of cervical tissues for CD4^+^ (F) and CD8^+^ T cells (G) using anti-CD4 labelled with FITC, anti-CD3 labelled with alexa 568 and anti-CD8 labelled with APC and observed under fluorescence microscope with 200X magnification. Inguinal lymph nodes collected from control and infected mice were minced to get single cell suspension and were treated with brefeldin A for 4hrs and stained for CD3, CD4 and CD8 and IFNγ. CD4^+^ and CD8^+^ T cells were analyzed for intra-cellular IFNγ expression in flow cytometer (Gating strategy is given in supplementary figure S2) (H). In a replicate experiment, CD4^+^ and CD8^+^ T cells were purified from lymph node single cell suspension and incubated for 48hrs to measure secreted IFNγ and TNFα levels by ELISA (I). Simultaneously part of the lymph nodes were lysed in TriZol and RNA was extracted to synthesize cDNA. *PD1, T-bet* and *CD40L* mRNA expression levels were measured using real-time qPCR (J). Splenocytes from spleen of infected and control mice were collected to purify CD4^+^ T cells and stimulated with *C*.*tr* lysate in presence of brefeldin A and analyzed for intracellular IFNγ and TGFβ expression using flow cytometry and plotted as % of cells (K) (Gating strategy was given in supplementary figure S2). (n=6; ***p<0.0001; **p<0.001)

### C. trachomatis induce anti-inflammatory phenotype and downregulate CD40 expression resulting in decreased T cell activation

*Chlamydia trachomatis* residing in the host cells can influence host machinery leading to changes favouring its survival and growth. Macrophages tend to polarise either to classical or alternatively-activated state based on external (T cell cytokines) or internal stimuli (metabolism or antigenic nature) [12, 13]. To elucidate the contribution of innate immune cells in mounting an immunosuppressive response observed during chronic chlamydial infection (Fig.1), we infected macrophages and epithelial cells with *C*.*tr*. Since, the outcome of the response would not only be dictated by the antigenic components but are also the result of changes in the host cells mediated by live bacteria (*C*.*tr*), heat killed bacteria with equal antigenic component (*C*.*tr*.Ag) was used to decipher the role of live bacteria mediated mechanism to alter innate immune cells. Macrophages primed with *C*.*tr*-Ag show increased expression of CD40, TLR4 and iNOS, while live *C*.*tr* primed macrophages show moderate levels of CD40 and increased arginase-1 expression (Fig.3A and D). This was concurrent with arginase activity and NO levels as well (Fig.3B and C). ROS levels were increased in both *C*.*tr* infection and *C*.*tr*-Ag priming, but they were higher in exposure to antigenic compared to live bacterial infection (Fig.3C). Macrophages primed with *C*.*tr*-Ag caused a significant increase in inflammatory cytokines and had decreased anti-inflammatory cytokines. However, macrophages infected with live bacteria demonstrated significantly lower levels of pro-inflammatory cytokines (TNFα and IL-12; except IL-1β), while increased levels of anti-inflammatory cytokine (IL-10 and TGFβ) was observed (Fig. 3E). To further analyse whether this alteration leads to differential polarisation of macrophages, we analysed the expression of M_1_ and M_2_ markers, MHC-II and CD206 respectively, in *C*.*tr*.Ag primed and *C*.*tr* infected macrophages. The macrophages primed with *C*.*tr*.Ag shows increase in cells with high MHC-II and low CD206 indicating the generation of M_1_-like macrophages, while the live bacteria skewed the macrophage phenotype to M_2_-like subset with low MHC-II and high CD206 expression (Fig.3F). Similar patterns of cytokine production and CD40 expression were observed when epithelial cells were primed with *C*.*tr*.Ag or live *C*.*tr* (Fig.3G and H). In summary, these results indicate that host cells mount an inflammatory immune response during priming with *C*.*tr*-antigens, however live *C*.*tr* may be manipulating the host machinery to its advantage by evading the essentialities required for bacterial clearance and causes phenotypic and functional skewing of macrophages.

**Figure 3:**
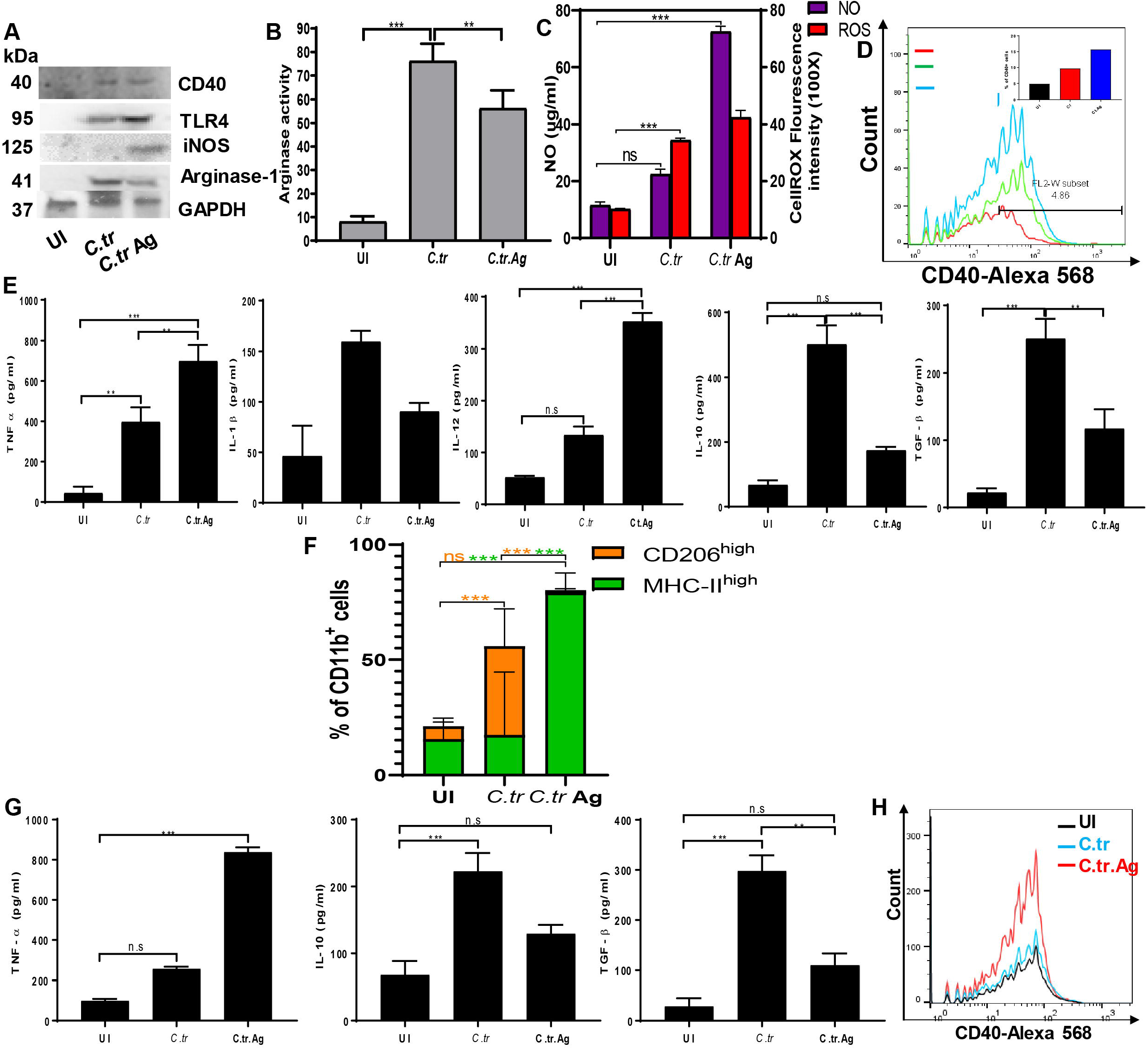
Live *C. trachomatis* induces immune suppression in innate immune cells. Bone marrow derived macrophages (BMDMs) were infected with *C*.*tr* or treated with *C*.*tr*-Ag for 18hrs and analyzed for CD40, TLR4, iNOS and Arginase-1 expression by western blot. GAPDH was used as loading control (A). In a replicate experiment, cells were lysed in PBS and arginase activity measured (B). BMDMs were treated with *C*.*tr* and C.tr-Ag for 12hr. NO levels were measured Griess reagent in supernatant. ROS levels were measured by CellROX deep Red in a Molecular devices fluorimeter (C). CD40 expression was analyzed on BMDMs in a replicate experiment by flow cytometer (D). After 24hrs of *C*.*tr* infection or C.tr-Ag treatment, BMDMs secreted cytokines (TNFα, IL-12, IL-10 and TGFβ) were measured in the supernatant using ELISA (E). *C*.*tr* infected or C.tr-Ag primed BMDMs were analyzed for CD206 and MHC-II expression to analyze macrophage polarization (F). (n=3; ***p<0.0001; **p<0.001). Mouse cervical epithelial cells were infected with *C*.*tr* or heat killed *C*.*tr*-Ag for 24 hrs and secreted cytokines (TNFα, IL-10 and TGFβ) were measured in the culture supernatant by ELISA (G). In a replicate experiment, CD40 expression was analyzed by flow cytometer (H).

To delineate the mechanism of immunosuppression and assess whether it is the contribution of *C*.*tr*-mediated cellular changes in macrophages or immunosuppressive T cells generated during the infection, we rechallenged the splenic T cells from all groups of mice by co-culturing them with differentially primed (*C*.*tr* or *C*.*tr*-Ag or left untreated) macrophages as explained above. The difference in T cell response from live *C. trachomatis* infected macrophages and *C*.*tr*.Ag primed macrophages might assist in delineating whether immune suppression is happening in macrophages or T cells. Following, the priming of macrophages and irradiating them, they were co-cultured with T cells harvested from spleen of infected and uninfected groups. CD4^+^ T cells from the acute infected mice show highest T cell proliferation and IL-2 secretion while the secondary reinfection group demonstrated low T cell proliferation and IL-2 production indicating decreased T cells activation (Fig.4A and B). Concurrently, IFNγ production was also decreased during chronic infection and secondary re-infection. Mice in the acute infection group, during later stages show high IL-4 indicating resolving phase, while the secondary infected group of mice secrete high TGFβ and IL-10 indicating an immunosuppressive response (Fig.4B). SI group of mice on secondary infection were not able to produce enough IFNγ. The T cells secreted cytokines from various groups were added to *C*.*tr* infected cells to analyse the anti-bacterial potential of secretome. The secretome form the acute infected mice rendered efficient clearance of the bacteria. However, the secretome of the chronic infected as well as secondary re-infection mice (C, CR and SI) failed to clear the bacteria. (Fig.4C). High TGFβ and IL-10 levels during chronic and secondary infections would have lead to immune suppression which might have resulted in poor bacterial clearance (Fig.1B-D). T cell activation and inhibitory receptors mRNA levels were analysed to understand the decrease in T cell activation and effector cytokine production. The decrease in T cells proliferation and functional immunosuppression could be attributed to increase in co-inhibitory receptors mRNA (*PD1, KLGR3, Tim3*) expression and decrease in *CD40L* mRNA levels in T cells indicating the possibility of anergic and exhaustive T cell population. The continued exposure of antigen during chronic infection would have led to increase in co-inhibitory molecule receptors mRNA (*PD1, KLGR3, Tim3*) expression. T cells from secondary infection also show increased in these markers with increased TGFβ secretion indicating that the T cells were inefficient and are of probably anergic and exhaustive phenotype (Fig.4D). Previously, high TGFβ at the site of infection has been reported to induce T cell anergy and exhaustion [14]. The proliferation and activation of CD8^+^ T cells was same as observed in CD4^+^ T cells (Fig.4E). CD8^+^ T cells from acute infection show higher IFNγ and TNFα secretion and expressed high *granzyme* and p*erforin B* mRNA levels compared to chronic and secondary infections (Fig.4F and G). Secondary infection after chronic infection was not able to elicit anti-chlamydial immune response, however macrophages primed with *C*.*tr*-Ag show better response compared to live bacterial infection (Fig.4). This indicates that live *C*.*tr* mediates refractiveness in innate immune cells rendering immune suppression via their cognate interactions or cytokines.

**Figure 4:**
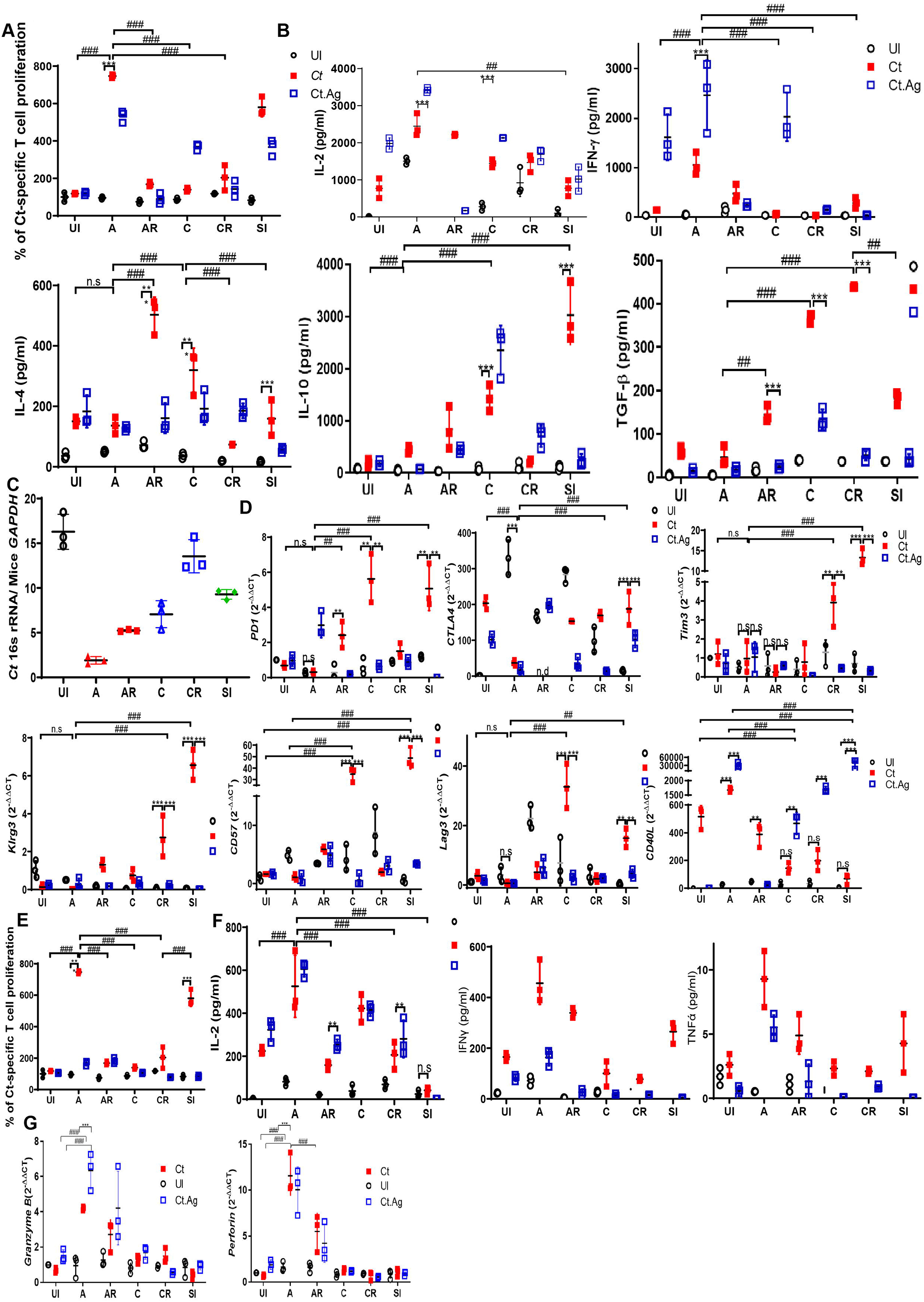
*C. trachomatis* chronic infection decapitates T cells. Splenocytes were isolated from spleen of infected and control mice. CD4^+^T cells isolated from splenocytes were co-cultured for 48hrs with fixed uninfected macrophages, *C*.*tr* infected macrophages and C.tr-Ag treated macrophages. T cell proliferation was analyzed by MTT assay (A). Secreted cytokines (IL-2, IL-10, IL-4, βγ) were measured from supernatants by ELISA (B). The supernatant from the *C*.*tr* infected macrophages-CD4^+^ T cells co-culture was added to the *C*.*tr* infected McCoy cells to analyze the effect of the secretome on bacterial viability. C.tr burden was measured as genomic ratio of *C*.*tr* to *Mus musculus* by real-time qPCR (C). In a parallel experiment co-cultured CD4^+^ T cells were collected in TriZol after 12hrs and mRNA expression was analyzed for *CD40L, PD1, CTLA4, Tim3, Klrg3, Lag3* genes using real-time qPCR and plotted as 2^-ΔΔCT^. GAPDH was used as control(D). CD8^+^ T cells isolated from splenocytes were co-cultured for 48hrs with fixed uninfected macrophages, *C*.*tr* infected macrophages and C.tr-Ag treated macrophages. CD8^+^ T cell proliferation was analyzed by MTT assay (E). In a parallel experiment, after incubation for 48hrs, secreted cytokines (IL-2, IFNγ and TNFα) were measured in the supernatant by ELISA (F). In a parallel experiment with CD8^+^ T cells, granzyme and perforin mRNA levels were measured by real-time qPCR (G). (n=6; ***p<0.0001; **p<0.001)

### CD40 activation potentiates anti-inflammatory immune response via regulation of macrophage activation

CD40 plays a major role in T cell activation and dictates the outcome of immune response by signalling bidirectionally to both innate and adaptive immune cells. CD40 expression levels and activation state has been shown to control inflammatory output and subsequent T cell differentiation and activation. *C*.*tr* manipulates CD40 expression on macrophage as well as epithelial cells as depicted earlier (Fig.3A, D and H). Thus, to understand the role of decreased CD40 expression by *C*.*tr* on inflammatory output, CD40 signalling was activated in infected cells using activating CD40 antibody (CD40^a^; clone: 5C3). CD40 activation results in activation of ERK1/2, p38 pathways to exert its effects. These pathways can also be activated by multiple upstream signals, as *C*.*tr* infection partially activate these pathways to various degree (Fig.5A). CD40 activation resulted in increased p38 phosphorylation and decreased ERK1/2 phosphorylation in uninfected macrophages. However, during *C*.*tr* infection a skewed response with substantial decrease in p38 activation and increase in ERK1/2 phosphorylation was observed. This effect is dose and time dependent (Fig.5B and C). CD40 activation mediated changes in infected cells resulted in significant decrease in IL-12 secretion, but TNFα and IL-1β secretion were comparable to the control (Fig. 5D). CD40 activation in infected macrophages, lead to increased secretion of anti-inflammatory cytokines IL-10 via ERK1/2 signalling (Fig. 5D). To understand the necessity of CD40-CD40L induced changes on T cells, uninfected and infected macrophages were co-cultured with CD4^+^ T cells in presence or absence of blocking CD40 antibody (CD40^b;^ clone: HM40-3). Blocking of CD40 on the APC lead to decrease in IFNγ and IL-4 secretion without altering the IL-17 and TGFβ secretion highlighting the importance of the CD40 on APCs to regulate the Th1/Th2 differentiation (Fig. 5E and F) [15, 16]. Further, infected macrophages were co-cultured with T cells expressing varying levels of CD40L (Th1-*CD40L*^high^; Th2-*CD40L*^med^; Treg-*CD40L*^low^) (SF.5A), T cells with higher CD40L expression were not able to increase IL-12 secretion significantly nor modulate IL-10 secretion, whereas T cells expressing low CD40 expression were able to induce IL-10 secretion from macrophages (Fig.5G). These indicates that decreased CD40 levels on *C*.*tr* infected macrophages plays a major role in translation of immune response to T cells inducing immunosuppression.

**Figure 5:**
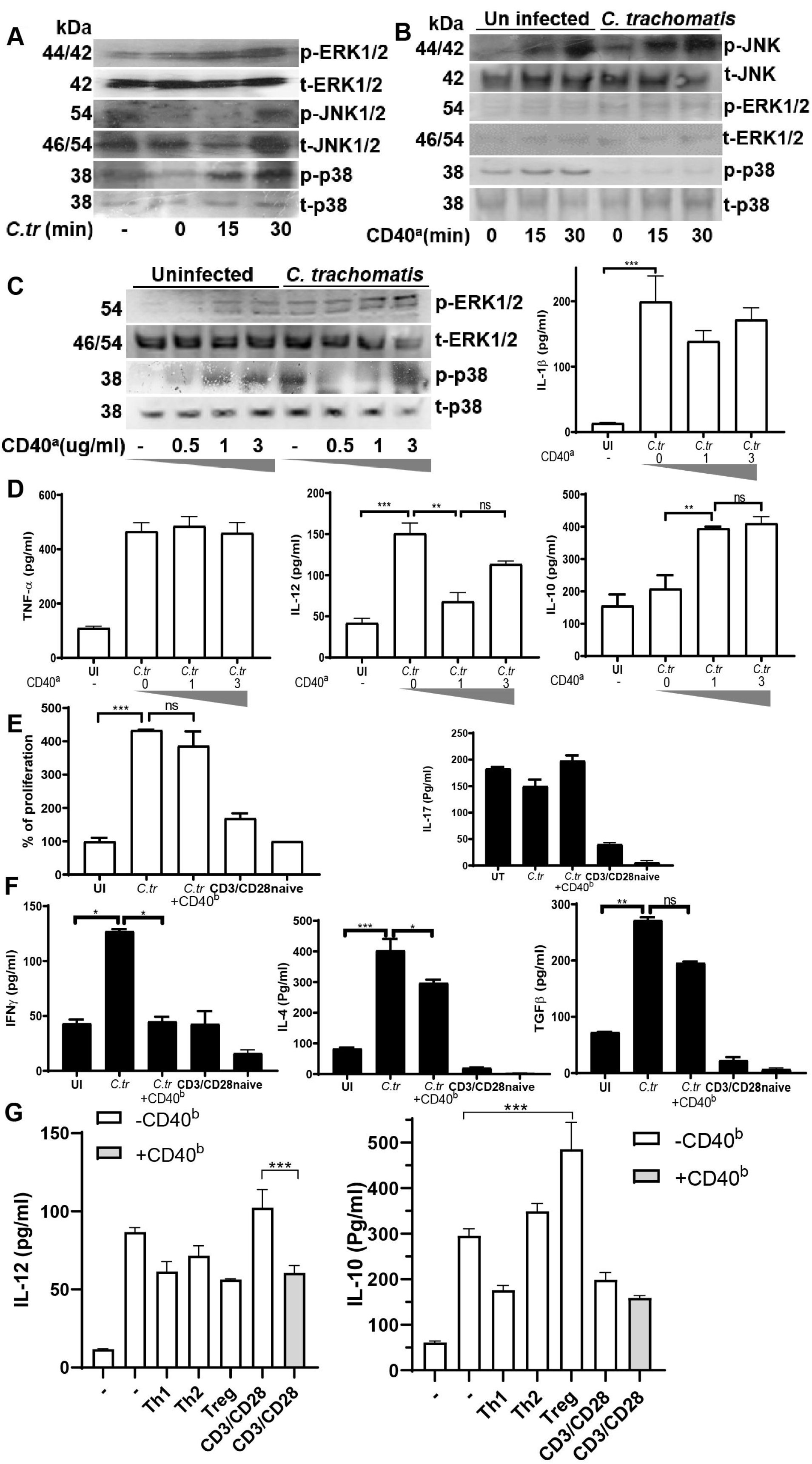
CD40 activation induces immune suppression during *C. trachomatis* infection. BMDMs were infected with *C. trachomatis* and collected at indicated time points. Total and phosphorylated EKR1/2, JNK1/2 and p38 proteins were analyzed by Immunoblotting (A). CD40 was activated for indicated time points in uninfected and *C*.*tr* infected cells with CD40 activating antibody (CD40^a^; clone:5C3) and analyzed for JNK, ERK and p38 pathway by immunoblotting (B). CD40 was activated on uninfected and *C*.*tr* infected BMDMs, with various doses of CD40 activating antibody and EKR1/2 and p38 pathway was analyzed by immunoblotting (C). CD40 was activated with anti-CD40^a^ on *C*.*tr* infected BMDMs for 24hr and analyzed for secreted cytokines using ELISA (D). *C*.*tr* infected BMDMs were co-cultured with CD4^+^ T cells in presence or absence of CD40 blocking antibody (CD40^b^; clone:HM40-3) and T cell proliferation was analyzed by MTT assay (E). In a parallel experiment, after 48hr of co-culture, cytokines (IFNγ, IL-4, IL-10 and TGFβ) were measured in the supernatant by ELISA (F). Naïve T cells obtained from Balb/c mice were differentiated to Th1, Th2 and Treg in presence of IL-12, IL-4; and TGFβ and retinoic acid respectively. In all groups CD3/CD28 and IL-2 were added to stimulate T cell activation. CD40L expression was analyzed (SF5). *C*.*tr* infected macrophages were co-cultured with fixed Th1 (CD40L^high^), Th2 (CD40L^int^), Treg (CD40L^low^) in presence or absence of CD40^b^ for 24hr. IL-12 and IL-10 levels measured by ELISA (G) (CD40L mRNA expression levels were given in SF5). (n=3; ***p<0.0001; **p<0.001)

### Low IFNγ mediates persistence of C. trachomatis which in turn polarize macrophages to M_2_ phenotype

Bacterial infection induce inflammation and activate T cells which depend on MHC-TCR, co-stimulation including CD40-CD40L and released cytokines. During intracellular infections, Th1 cytokines potentiates the clearance of bacteria, while Th2 cytokines participate in achieving the resolution phase. During intracellular infections, T cell secreted IFNγ facilitates the clearance of bacteria by various mechanisms including but not limited to (i) enhancing antigen presentation by MHC and (ii) depleting tryptophan by IDO [17, 18]. *C. trachomatis* decreases CD40 resulting in decreased T cell activation which in turn results in decreased IFNγ production, thus evading the host immunity. To understand the role of IFNγ on *C*.*tr* survival intracellularly, various doses of IFNγ was added for different time points to infected cells. 10ng/ml of IFNγ cleared the bacteria in 12hrs. No intracellular *C*.*tr* was observed even after removal of IFNγ indicating the complete clearance of *Ctr*. 5ng/ml of IFNγ cleared intracellular *C*.*tr* but complete clearance was not observed even after 24hrs of IFNγ exposure. 1ng/ml of IFNγ decreased the bacterial burden, but the removal of IFNγ resulted in increased presence of *C*.*tr* (Fig.6A). This can be compared to natural infection of *Chlamydia trachomatis* where acute infection accompanies high IFNγ (as shown in Fig.1G and J), while low IFNγ groups can be compared to persistent infection. Belland et al., have also reported that insufficiency of IFNγ leads to aberrant form of *C*.*tr* which mediates persistence and subsequent progression to latent infection. The latent infection with *C*.*tr* has been reported to progress to fallopian tube damage (Vicenfeld Xe et al., 2002, J. Infectious diseases). The infected cells exposed to high IFNγ could not reinfect the fresh batch of cells while infected cells exposed to low IFNγ were able to infect fresh cells (Fig.6A). The later failed to release TNFα, demonstrated high IL-10 secretion and failed to increase CD40 expression compared to *C*.*tr* infected cells with no or low IFNγ treatment (Fig.6B). This may be due to generation of aberrant form of *C*.*tr* in presence of low IFNγ compared to the normal *Ct* infection. IFNγ is known to induce indoleamine 2,3-dioxygenase (IDO) activity and nitric oxide synthesis facilitating pathogen clearance. However, *C*.*tr* infection prevented an increase in IDO activity and NO levels on IFNγ addition (Fig.6C and D). Additionally, aberrant *Ct* infection failed to increase CD40 expression on epithelial cells and macrophages (Fig. 6E and Fig.7A). IFNγ was able to induce CD40 expression on infected epithelial cells at higher concentration but it reverted back to the original levels following the removal of the cytokine (Fig.6E). The results indicate that the high IFNγ levels secreted from T cells during infection might clear bacteria and induce inflammation, but lower levels or absence of IFNγ during infection causes immunosuppression. Thus, absence or low levels of IFNγ can cause more damage to the host than the infection alone. These observations were also found in chronic infections of *C. trachomatis* earlier by *[19]*. Infected epithelial cells secretome could drive macrophages to express *CD40, CD206* and *arginase-1* mRNA indicating alternatively activated macrophages. IFNγ addition to epithelial cells did not show significant change in the macrophage activation via their secretome (Fig. 6F). Previous reports also indicate that the presence of alternatively activated macrophages or Th2 at the site of chlamydial infection can contribute to immunosuppression. [19, 20]. Similarly, epithelial cells might also be the culprit for the immune suppression as indicated (Fig.6).

**Figure 6:**
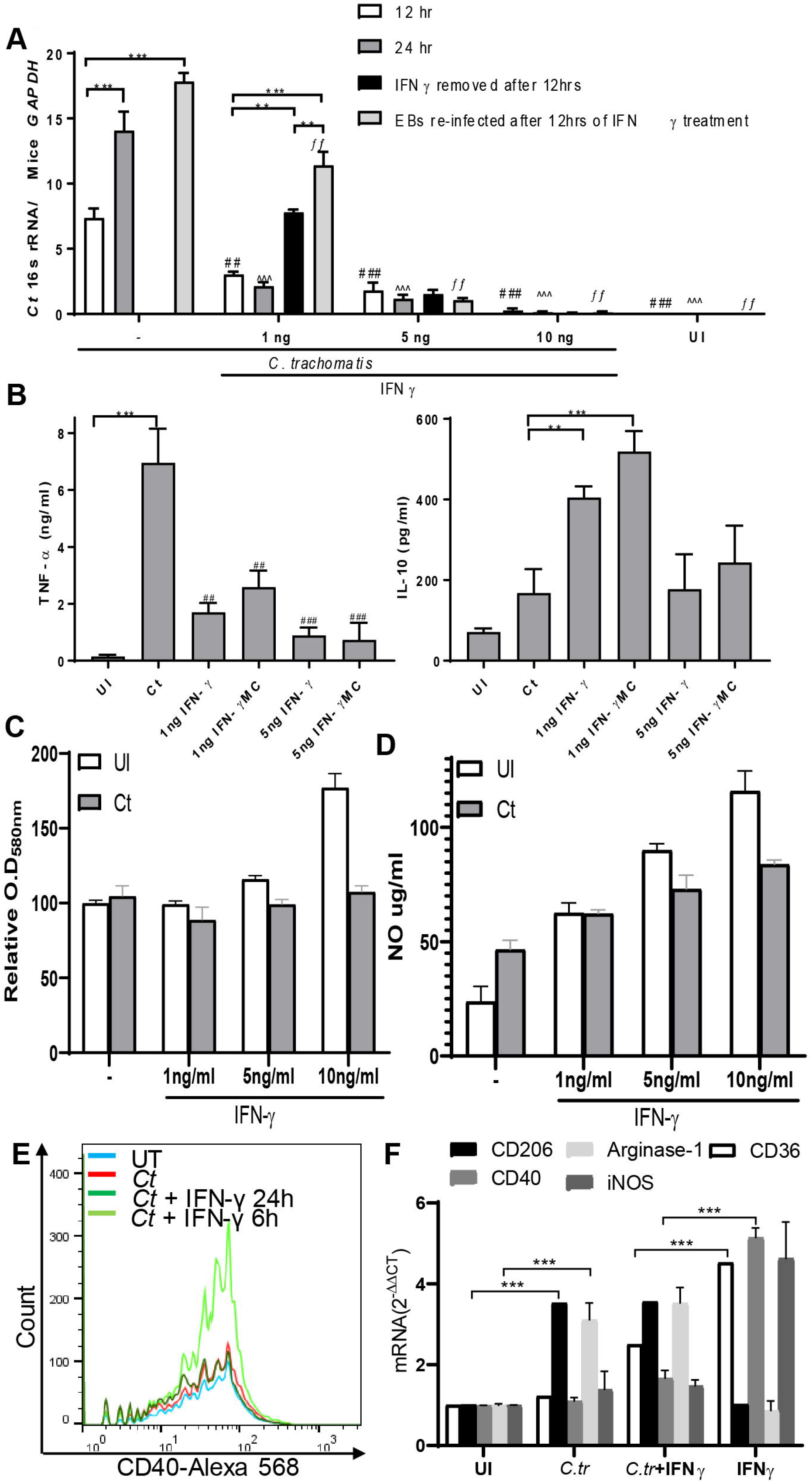
IFNγ insufficiency leads to incomplete clearance of *C*.*tr* in epithelial cells. Mouse epithelial cells were infected *C*.*tr* and treated with various doses of IFNγ for 12hr and 24hrs. To replicate the IFNγ unavailability, IFNγ was removed after 12hrs and incubated for 12 hr more (black bar). The *C*.*tr* collected from the replicate well were re-infected to analyze their ability to infect fresh cells (grey bar). Bacterial burden was analysed by real-time qPCR (A). Epithelial cells infected with *C*.*tr* were treated with low (1ng/ml) and medium (5ng/ml) IFNγ and analyzed for TNFα and IL-10 by ELISA (B). In parallel experiments, IDO activity (C) and NO levels (D) were measured after 24hrs. CD40 levels were measured on mouse cervical epithelial using flow cytometer (E). The secretome from the infected cells in presence or absence of IFNγ(5ng/ml) were added to naïve macrophages and analyzed for mRNA expression of *CD36, CD206, iNOS, CD40* and *arginase-1*. Naïve macrophages and equal amount of IFNγ supplementation served as negative and positive control (F). (n=3; ***p<0.0001; **p<0.001)

**Figure 7:**
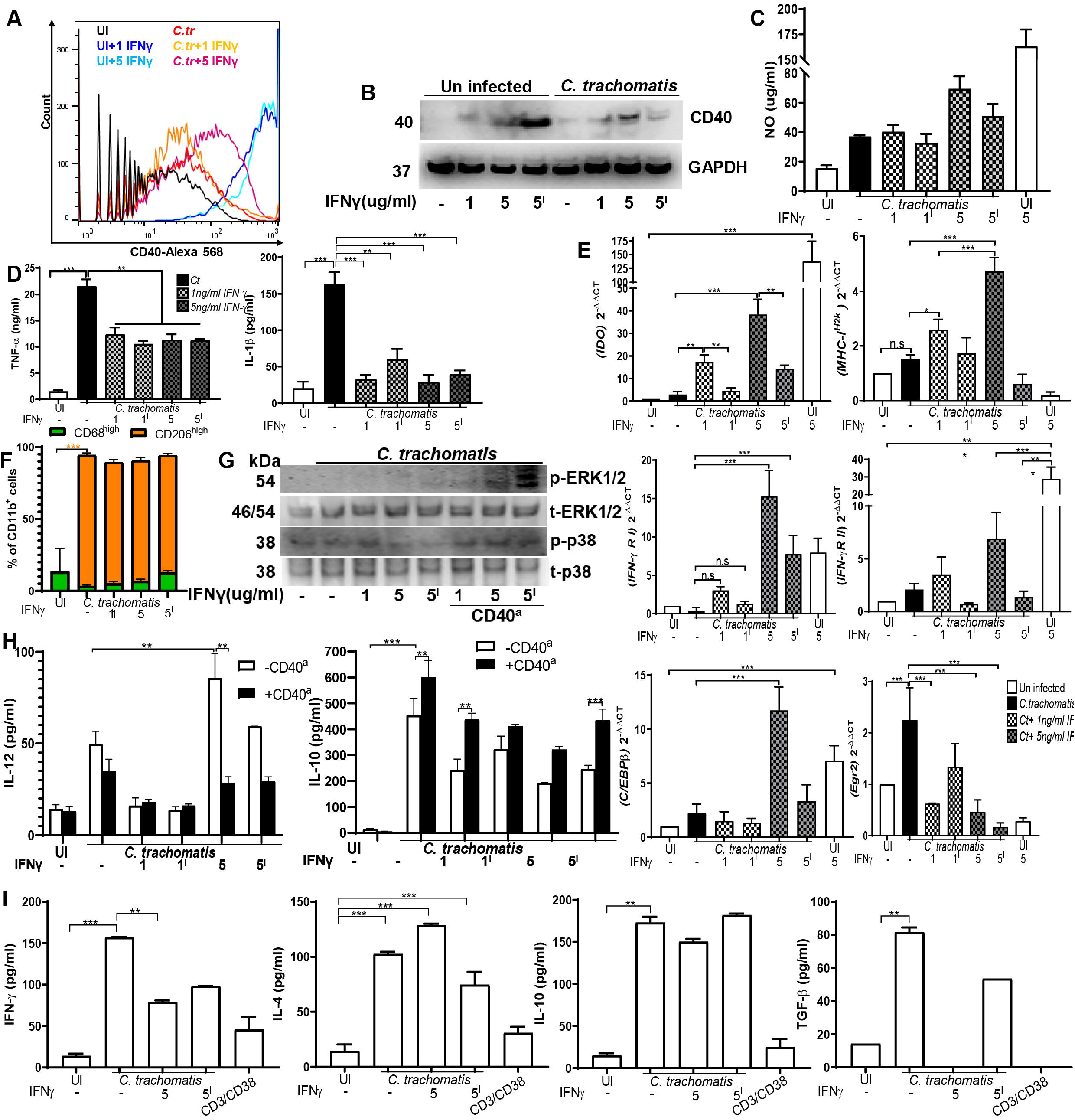
IFNγ insufficiency during *C*.*tr* infection cannot induce inflammatory response. *C*.*tr* infected with BMDMs were treated with 5ng/ml IFNγ either for 24hr or 6hr and analyzed for CD40 expression (A). In a parallel experiment, CD40 levels were measured by immunoblotting (B). *C*.*tr* infected BMDMs were treated with 1ng/ml or 5ng/ml of IFNγ. IFNγ was removed after half the time and replaced with fresh media (^I^). After 24hrs, NO levels were measured by Griess reagent (C). TNFα and IL-1β levels were measured by ELISA (D). In a replicate experiment, after 8hrs, mRNA levels of *IDO, MHC-I*^*H2k*^, *IFNγRI, IFNγRII, C/EBPβ* and *Erg2* were measured by real time-qPCR. GAPDH was used as control (E). BMDMs were infected *C*.*tr* and exposed to varying concentrations of IFNγ after 6hrs of infection (1 and 5) and replace with fresh media after 12hrs of IFNγ supplementation (5^I^). The cells were analyzed for CD11b, CD68 and CD206 using anti-CD11b labelled with APC, anti-CD68 labelled with FITC and anti-CD206 labelled with PE in flow cytometer for macrophage polarization ad expressed as % of CD11b^+^cells (F) (Gating strategy was given in SF6A-B). CD40 was activated on *C*.*tr* infected BMDMs supplemented with IFNγ as mentioned above with CD40^a^ for 30min and analyzed for total and phosphorylated EKR1/2 and p38 (G). In a similar experiment, CD40 was activated for 18hrs, secreted IL-12 and IL-10 were measured by ELISA (H). *C*.*tr* infected macrophages treated with 5ng/ml either for 24hr or 12hr were co-cultured with Naïve T cells for 48hrs. Secreted T cell cytokines (IFNγ, IL-4, IL-10 and TGFβ) were measured by ELISA from supernatant. Naïve T cells treated with activating CD3 and CD38 antibodies were used as positive control (I). (n=3; ***p<0.0001; **p<0.001)

Macrophages are critical for the clearance of bacteria and influence the micro-environment. *C*.*tr* infected macrophages show increased CD40 expression moderately, while the expression is increased in the presence of high IFNγ only, not at lower concentration (Fig.7A and B). iNOS, the downstream regulator of the IFNγ produces NO and was increased in IFNγ exposed uninfected macrophages, but not in *C*.*tr* infected macrophages (Fig.7C). Even though IFNγ could increase the CD40 expression moderately, it resulted in decrease release of TNFα and IL-1β significantly (Fig.7D). Macrophages infected with *C*.*tr* show decreased expression of *MHC-I*^*h2k*^, *IFNγRI* and *IFNγRII* mRNA levels poor indicating antigen presentation and failed immune activation. IFNγ increased *IDO* expression in both infected and un-infected macrophages, but infected cells exposed to IFNγ show decreased *IDO* mRNA levels compared to IFNγ treated un-infected cells. This indicates mechanism employed by *C*.*tr* against host immune mechanisms. Exposure of macrophages to low levels of IFNγ increased *MHC-I*^*h2k*^ but not *IFNγRI* and *IFNγRII*, however exposure with high levels of IFNγ enhanced *IFNγRII* but the removal of IFNγ resulted in decreased *IFNγRII* expression (Fig.7E). Egr2, transcription factor regulating the transcription of immunosuppressive genes [21] was decreased following treatment with high IFNγ, while *C/EBPβ* a transcription factor for inflammatory genes was unaltered. *Egr2* and *CEBP/β* levels indicate that IFNγ can modulate macrophage activation state in infected cells transiently (Fig. 7E). Flow cytometric analysis show that IFNγ supplementation was not able to skew infected macrophages from alternative activated state (expressing CD206) to classically activated macrophages (CD36 expression) (Fig. 7F). As, IFNγ was not able to increase the CD40 expression significantly and skew macrophages to M_1_, we analysed the role of CD40 activation on IL-10 and IL-12 cytokines. CD40 activation even following IFNγ addition leads to ERK1/2 activation in macrophages leading to increased IL-10 secretion (Fig.7G and H). While IFNγ addition on uninfected macrophages increases CD40 expression (Fig.7A) and subsequent increase in p38 activaton (Fig.7G) resulting in IL-12 secretion (Fig.7H). CD4^+^ T cells co-cultured with aberrantly infected macrophages revealed that T cells activation was low as indicated by proliferation (Supplementary Figure SF). Concurrently, T cells also secreted low IFNγ during aberrant infection (^I^) compared to normal infection. No significant change was observed in IL-10 and IL-4 release indicating that persistence infection does not alter immunosuppressive T cells (Fig. 7I). IFNγ mediated persistent infection in macrophages were able to skew T cell differentiation to TGFβ producing T cells (Fig. 7I). These results evidently show that, low IFNγ mediated persistent infection skews macrophages to alternatively-activated macrophages (M_2_) and the prime the T cells to anti-inflammatory phenotype there by resulting in an immune-suppressive response.

### M_2_ macrophages demonstrate higher bacterial burden and mount an anti-inflammatory response

Infected macrophages demonstrate M_2_ phenotype, similarly the supernatant of infected epithelial cells can also skew macrophages to M_2_ phenotype. To understand the role macrophage polarisation and their effect on infection status, polarised macrophages were infected with *C*.*tr*. M_2_ macrophages harbour high bacterial burden compared to M_1_ subset (Fig.8A). Previously, I Tietzel et al., have shown that alternatively activated macrophages harbour more bacteria compared to classically activated macrophages (31134002). M_2_ macrophages on *C*.*tr* infection increased the expression of co-inhibitory molecule *PDL-1* and decreased *CD68* mRNA levels along with decreases *iNOS* expression (Fig.8B). M_2_ macrophages show increased expression of *protein disulfide-isomarese* (*PDI)* and *Erphin B2*, which are responsible for *C*.*tr* binding, which makes macrophages more suitable to higher *C*.*tr* burden (Fig. 8B). Infected macrophages have high *Egr2* expression, while uninfected M_1_ macrophages alone show high *C/EBPβ* mRNA. CD40 expression also decreased on M_1_ and M_2_ macrophages (Fig.8C). M_1_ macrophages on infection were skewed towards M_2_ phenotype expressing higher CD206 levels compared to uninfected (Fig.8D). M_1_ macrophages also increased arginase activity on *C*.*tr* infection indicating skewing of macrophages to M_2_ phenotype (Fig.8E). M_2_ macrophages on infection secreted high levels of signature cytokine IL-10 (Fig.8F). Infection of the M_1_ macrophages with *C*.*tr* also lead to secretion of IL-10. Classically-activated macrophages during *C*.*tr* infection secrete inflammatory cytokines: TNFα and IL-12 but also accompanies IL-10 secretion (Fig.8F). Macrophages show plasticity based on the cytokine availability. M_2_ infected macrophages were exposed to varying concentrations of IFNγ to analyse to ability to revert M_2_ to M_1_. IFNγ was not able to increase CD40 expression on infected M_2_ macrophages and retain its polarisation, even after addition of IFNγ which might be due to decreased availability of its receptor (Fig.8G-I). CD40 activation in infected polarised macrophages resulted in significant increase of IL-10 but not IL-12 (Fig.8J). To analyse the effect of the polarised macrophages infected with *C*.*tr* on CD4^+^ T cells, infected macrophages were co-cultured with naïve CD4^+^T cells. In a macrophage-T cell co-culture the M_0_ macrophages induce T cell proliferation and accompanies the release of the release of INF-γ, IL-4 and IL-10. While M_2_ macrophage-T cell co-culture demonstrate poor proliferation and enhanced levels of IL-10 and IL-4 and significantly lower levels of IFNγ. This indicates that M_2_ subset prime anti-inflammatory immunosuppressive response in T cell co-culture, however, the T cell co-cultured with infected M_1_ macrophages did not have increased released of IL-10, IL-4 and TGFβ, but they were handicapped in their potential to release IFNγ (Fig.8K). The CD40-CD40L interactions forms a bridge between innate and adaptive immune cells moderating the immune response. In order to examine the decreased CD40 expression and M_2_ polarisation during *C*.*tr* infection macrophages on adaptive immune cells, infected polarised macrophages were co-cultured with B cells. It is evident that M_2_ macrophages were able to induce higher B cell proliferation compared to M_0_ macrophages and the effect is abrogated on CD40 blockade indicate its role in mounting effective adaptive immune response (Fig.8L). In overall, *C*.*tr* skews macrophages to M_2_ and hinder CD40 expression, thus incapacitating T cell activation for prolonger survival.

**Figure 8:**
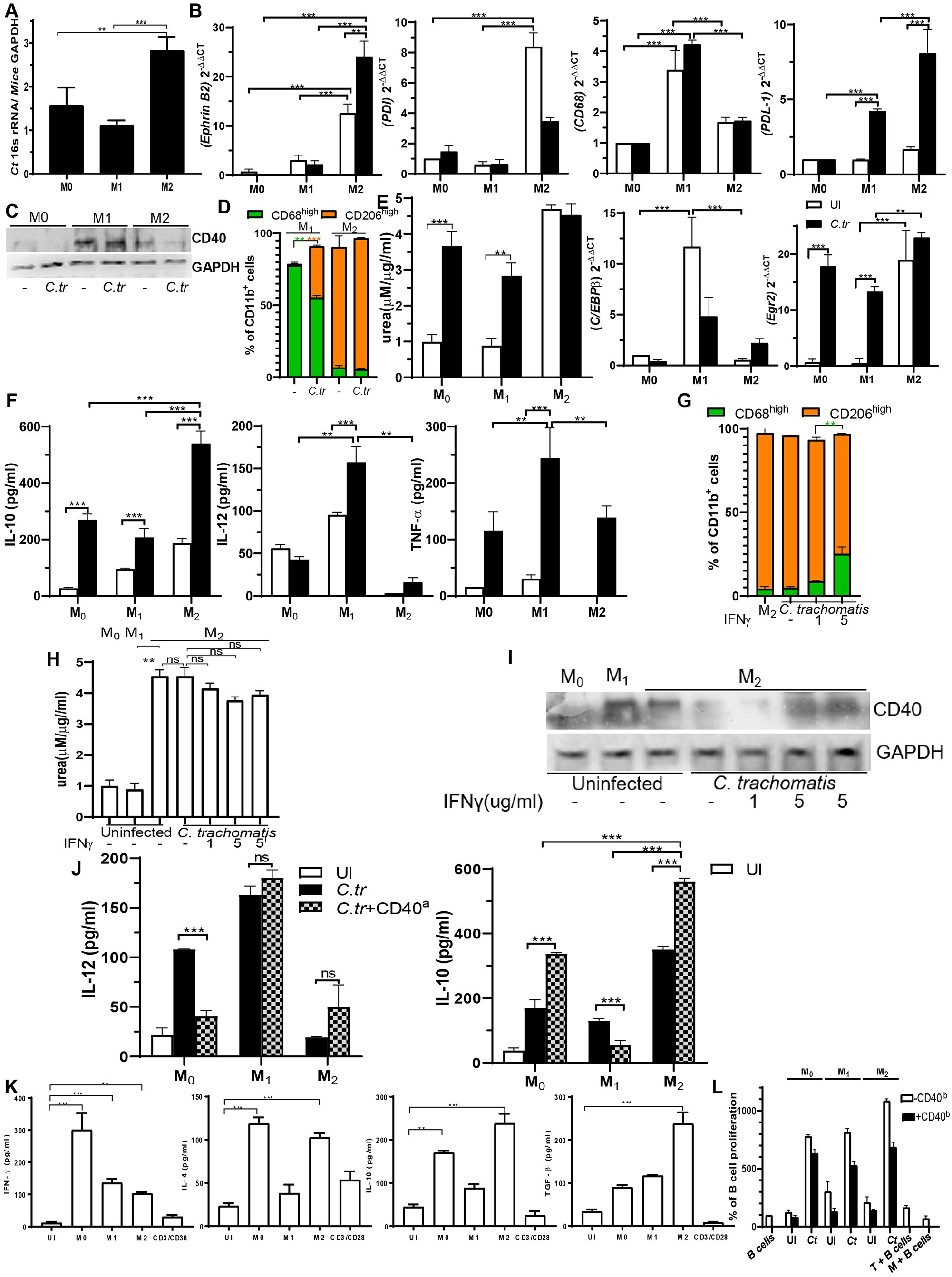
M_2_ macrophages harbor higher *C*.*tr* and induces immune suppression. Bone marrow derived cells were differentiated to M_0_(M-CSF), M_1_(GM-CSF and IFNγ) and M_2_(M-CSF, IL-4 and IL-10). Polarized macrophages were infected with *C*.*tr* and analyzed for bacterial burden using real time qPCR measuring *C*.*tr* and *Mus musculus* genome (A). Polarized macrophages were infected with *C*.*tr* for 6hrs and mRNA levels of indicated genes were analyzed by real-time qPCR (B). CD40 expression were analyzed by immunoblotting after 18hrs of infection (C). M_1_ and M_2_ polarized macrophages were infected with *C*.*tr* for 24 hrs and analyzed for CD11b, CD206 and CD68 expression using flow cytometer and represented as % of CD11b^+^cells expressing CD68^high^(M_1_) and CD206^high^(M_2_) (D) (Gating strategy was given in SF6A and C). Arginase activity was measured in polarized macrophages after 24hrs of infection (E). IL-10, IL-12 and TNFα secretion from polarized macrophages infected with *C*.*tr* were measured from supernatant after 24hrs by ELISA (F). M_2_ macrophages infected with *C*.*tr* were treated with indicated concentration of IFNγ and analyzed for macrophage polarization by measuring the expression of CD11b, CD206 and CD68 in flow cytometry (G) (Gating strategy was given in SF6A and D). Similarly, arginase activity was measured in lysates from replicate experiment (H). In a parallel experiment, CD40 expression was analyzed by immunoblotting after 18hrs (I). CD40 was activated on *C*.*tr* infected polarized macrophages using CD40^a^ for 24hrs and analyzed for secreted IL-10 and IL-12 levels by ELISA (J). Polarized macrophages infected with *C*.*tr* were co-cultured with naïve T cells and analyzed for secreted T cell cytokines (IFNγ, IL-4, IL-10 and TGFβ) from supernatant after 48hrs (K). The above experiment was repeated in presence or absence of CD40 blocking antibody added to the macrophages before co-culturing with T cells. After 24hrs of co-culture, naïve B cells were added to the co-culture and B cell proliferation was analyzed by MTT assay (and CD19 expression as given in supplementary figure S). T cells, B cells without macrophages; macrophages and B cells without T cells and Naïve B cells served as negative controls (L). (n=3; ***p<0.0001; **p<0.001)

## Discussion

Latent *C. trachomatis* infection in human is reported to cause of infertility and ectopic pregnancy. Previously, researchers have been using *C. muridarum* -genital infection or *C. pneumoniae-*lung infection model to understand and mimic the immune response, immunopathology and immunosuppressive mechanisms of *C. trachomatis* at mucosal site. [22]. However, the molecular mechanism of the immunosuppression is still unclear. In this study, we sought to elucidate the immunosuppressive innate immune mechanisms employed by *C. trachomatis* that contribute to the generation of defective T cell response leading to persistent infection.

We have used mice model for acute, chronic and repeated infections for *C. trachomatis* as explained in the schematic (Fig.1A). We found that bacterial shedding stops 45 days post infection, but the *C*.*tr* was able to reside in the cervical tissue at lower numbers. To understand the immunotolerance, Balb/c mice infected on day 60 were observed for bacterial burden and analyzed for various immune parameters. During primary infection, *C. trachomatis* stimulates IFNγ production from both CD4^+^ T cells and CD8^+^ T cells that subsequently controls the its replication and clears infection [23]. However, upon rechallenge, the response of these cells is significantly lowered compared to the primary infection, with fewer CD4^+^ T and CD8^+^ T cells (Fig.2-H). This impaired secondary immune response during chronic infection exhibits an exhausted phenotype defined by decreased proliferation, lower effector cytokine production. It also demonstrated markers of T cell anergy and exhaustion. This might contribute to persistent infection as indicated by extended presence of bacteria in genital tract. Low IFNγ levels following poor activation of the T cells or low IFNγ producing T cells also induce aberrant form of *C. trachomatis* [24]. This might be due to the presence of M2 macrophages at the cervix during infection. The infected macrophages and epithelial cells show poor CD40 expression which is required for inducing Th1 activation (Fig.6E and 7A-B). Immunosuppressive cytokines (TGFβ and IL-10) secreted at the cervix have earlier been reported to increase co-inhibitory molecule expression and cause immune suppression [25]. Similarly, we report that *C*.*tr* infection resulted in poor T cell activation, decreased *CD40L* mRNA expression and increase *PD1, Klrg3* and *Ctla4* mRNA expression. This failure in T cell activation might be contributing to persistent infection of *C*.*tr*.

To delineate the mechanism of immunosuppression during chronic infection, macrophages and epithelial cells were infected with C.*tr* Infected cells demonstrated enhanced immunosuppressive cytokines (IL-10, TGFβ) (Fig.3). *C*.*tr*-Ag used as control, demonstrated increased release of inflammatory cytokines, enhanced CD40 expression and polarised macrophages to M_1_ phenotype indicating that, may be only live bacteria have the capability to modulate immune response via invasion and viability associated factors which warrants separate investigation (Fig.3). Previously, chlamydial CPAF, other effector proteins like CT868, CT166 and CT694 have been attributed to manipulate host cellular mechanisms [26, 27]. During intracellular infections, Th1 mediated bacterial clearance plays a major role in controlling the infection via generation of effector CD8^+^ T cells. The effective T cell activation happens via two stages, where T cell is primed via antigen presentation and secondary cognate interactions along with cytokines dictating the fate of T cells. However, during the secondary interactions, T cells are also able to skew the innate immune cell inflammatory output by regulating CD40 mediated cytokine release via CD40-CD40L interactions. CD40 levels were poorly increased during *C*.*tr* infection compared to its antigen counterpart, thereby leading to increased IL-10 on CD40 activation which was also reported in *Leishmania, E*.*coli, Bacteroides vulgatus* infections (10.1186/1471-2172-13-22) [28]. However, the CD40 levels are necessary for the induction of both Th1 and Th2 induction response (Fig. 3A and D), so poor CD40 expression on macrophages during chlamydial infection could be responsible for poor T cell activation in the tissue of infected mice (Fig. 1).

*C. trachomatis* infection stimulates IFNγ production from both CD4^+^ and CD8^+^ T cells, during primary infection, that subsequently limits its replication and clearance [23]. However, upon rechallenge the response of these cells is significantly lowered, with fewer CD4^+^ and CD8^+^ T cells. This impaired secondary immune response during chronic infection exhibit an exhausted phenotype defined by low effector expansion and cytokine production with increased T cell anergy and exhaustion markers (Fig.4D). This leads to persistent infection as indicated by continued presence of bacteria in the genital tract. Low IFNγ levels resulting from decreased activation and fewer IFNγ producing cells also induce aberrant form of *C. trachomatis* [29, 30]. Upon removal of IFNγ from cultures or decreased CD4^+^IFNγ^+^Th1 cells, *C. trachomatis* replicates and leads to sustained infection as indicated by bacterial burden. These persistently infected macrophages also show anti-inflammatory properties (Fig.7F and H). Further, M_2_-like macrophages also contribute to increased immunosuppressive cytokine, TGFβ and IL-10, which can derail the activation of IFNγ producing T cell activation and sustain IL-4, IL-10 and TGFβ producing T cell activation contributing to impaired bacterial clearance.

Our data provides a possible mechanism by which the T cell response is impaired during *C. trachomatis* infection. During primary acute infection with *C. trachomatis*, it is evident that macrophages are active and translates the inflammatory response to activate T cell in mediating IFNγ production. On repeated infection, macrophages and epithelial cells decreases CD40 expression limiting T cell activation. Macrophages during chronic infection are skewed to M_2_ phenotype that release increased IL-10 and TGFβ. These cytokines supress T cell activation in addition to PDL1/PD1 mediated co-inhibitory interaction. IFNγ restricts *C. trachomatis* growth, and hence lack of this cytokine during chronic infection makes mice more susceptible to infection. This is consistent with our data demonstrating increased bacterial burden during chronic and latent infection. Previous reports have demonstrated high IL-10 and TGFβ production in human cervical tissue following *C*.*tr* infections [31, 32]. PDL1 expression during *C. trachomatis* infection was also reported to be a culprit for reduced number of IFNγ producing CD8^+^ T cells that was attributed to incomplete bacterial clearance.

Importantly, our data show that differential macrophage polarisation during *C. trachomatis* infection is one of the key contributors to immunosuppression that primes the generation of exhaustive/anergic T cells. Although, macrophages are migrated to genital tract upon infection, the major difference between acute and chronic infections is functional difference with varied expression of CD40 mediating differential regulation of T cell. Our data provide interesting evidence of C.*tr* regulation of macrophage polarisation which could explain, in part, T cell insufficiency during chlamydial infection. Previous reports also indicate that high PDL1 expression contribute to decreased T cell proliferation and effector cytokine production from T cells. Our results indicate that this regulation is dictated by the cells of innate immune response. Therapeutic strategies to control *C. trachomatis* infection and promote effector and memory T cells to efficiently clear the *C*.*tr* infection should focus on preventing the generation of M_2_ macrophage or reverting the Chlamydia-mediated M_2_ macrophage polarisation. Thus, an attractive strategy to regulate *C*.*tr* infection would be to focus on altering the cellular and molecular innate immune mechanisms that will subsequently orchestrate the improved effector T cell function. However, these provide very early indication on the role of polarised macrophages in dictating the persistence/clearance of *C*.*tr*. However, a detailed study is warranted to establish the mechanism of macrophage polarisation by *Chlamydia trachomatis*.

## Materials and Methods

### Reagents

All the recombinant proteins and antibodies used in the study are given in supplementary table 1 and 2 with details of dilution used, clone and brand.

### Mice

Age-matched Female BALB/c mice (6-8 wk old) were housed under specific pathogen-free conditions at the Animal house, Nirma university. All experiments were conducted in accordance with the guidelines of the Institutional Animal Ethical Committee after obtaining a clearance. Housing and handling of mice was in accordance with Committee for the Purpose of Control and Supervision of Experiments on Animals (CPCSEA).

Mice were infected with 10^6^ IFU (inclusion forming units) intravaginally in 100ul of SPG buffer repeatedly as shown in figure 1 for chronic (C) and chronic resolved (CR) infection for 20 days. Chronic infection group of mice were sacrificed on day 20 and CR mice were sacrificed on day 60. In a replicate group of chronic mice (C), secondary infection (SI) of *C. trachomatis* was done on day 60 and sacrificed on day 65. Acute infection (A) of *C. trachomatis* was done by infecting mice once and sacrificed on day 3. In a replicate group of A, mice were housed till day 20 for infection to resolve and termed as acute resolved group (AR). Control mice (UI) were treated with 4SP buffer intravaginally. Bacterial shedding and infection were monitored for the presence of bacteria in cervical swabs. Bacterial burden on the day of sacrifice was analysed in tissue and vaginal lavage (collected 1ml PBS).

### Bacteria and Bacterial lysate

*Chlamydia trachomatis serovar D* was procured from ATCC (VR-88). Chlamydia was cultured as previously described [33]. Briefly, frozen vials were thawed and inoculated to over-night grown confluent McCoy cells and observed for inclusion body formation. Elementary bodies (EBs) were purified from cell lysates of 48 hrs culture by 20%-50% Urograffin gradient centrifugation and stored in 4SP (0.4 M sucrose, 16 mM Na2HPO4) buffer at -80°C till further use. Inclusion forming units (IFU) of every batch was monitored before the experimentation. IFU vs Ct DNA 16S rRNA ΔCt values were normalised and used to determine the MOI during infections. 10^7^ *C. trachomatis* EBs were lysed in 0.1% tween at 70°C for 20min to prepare *C*.*tr* lysate which was later used as antigen (*C*.*tr*-Ag).

### Cells and cell culture medium

McCoy cell line was procured from NCCS, Pune, India and maintained in McCoy 5A media with 10% fetal bovine serum (FBS) at 37°C in 5% CO_2_.

Mouse Bone marrow derived macrophages (BMDMs) were generated as described elsewhere (Shah D et al., 2021). Briefly, bone marrow cells were collected from femurs of BALB/c mice and cultured in RMPI-1650 supplemented with 10% FBS. Cells were cultured in recombinant proteins M-CSF and GM-CSF for three days to generate macrophages M_0_/M_2_ and M_1_ respectively. Macrophages were further driven to M_1_ with IFNγ and M_2_ with IL-4 and IL-10 supplementation for three more days. On Day 6 macrophages (purity >95%) were washed with PBS before further experimentation (Supplementary table. 1).

T cells were purified from RBC-lysed splenocytes using nylon wool column [34]. CD4^+^T cells were purified from RBC-lysed splenocytes using Untouched CD4^+^ T cell enrichment kit (eBiosiences) using manufactures protocol.

Single cell suspensions from inguinal lymph nodes (ILN) were prepared by mincing the tissue between frosted end slides and the cells were used for flow cytometric analysis of surface and intracellular markers of T cells.

Mouse genital epithelial cells were isolated from female genital tract. Female genital tract including uterine horns were cut into small pieces after opening lumen and cleaning the mucus with PBS with 10µg/ml gentamycin. Small pieces of tissue were digested using 1% collagenase, 1% Tryspin and DNAse for 1hr at 37°C. Cells were collected from this mixture in DMEM with 5% FBS supplemented with Insulin and EGF. Cells were maintained by changing media every 2 days for 6 days and were harvested for further experimental procedures.

### *C. trachomatis* infection

EBs are added to cells at a MOI of 1 and centrifuged at 750g for 30min. After centrifugation, media was replaced with fresh media without antibiotics. Infected cells along with controls (4SP buffer only) were incubated for indicated time points.

### CD40 stimulation or blocking

CD40 activation on either infected or uninfected macrophages were done using CD40 activating antibody (CD40^a^; clone:5C3) after 30min of infection. CD40 blocking was done on macrophages 30min prior to addition of T cells in macrophage-T cell co-culture using CD40 blocking antibody (CD40^b^; clone:HM40-3)

### T cell differentiation assay

Naïve total T cells/CD4^+^T cells isolated from splenocytes were co-cultured with *C*.*tr* infected APCs/ C.tr-Ag primed APC/ uninfected APCs (APCs were fixed and washed with PBS). Naïve T cells without any stimulation served as a negative control. T cells were stimulated with CD3 and CD28 antibodies to analyse non-specific activation. After 3 days of co-culture, IFNγ, IL-10, IL-4 and TGFβ were quantified from the supernatant by ELISA. Simultaneously, in a replicate experiment, brefeldin A was added to co-culture after 6hrs and incubated for 48hrs to analyse intracellular IFNγ and TGFβ by flowcytometry.

### DNA isolation and qPCR for bacterial burden

DNA was isolated from infected cells as described earlier [35]. Briefly, cells were lysed in 10%SDS in TE buffer containing RNAse and Proteinase K. Phenol-chloroform method was used for DNA extraction. Extracted DNA was used as a template for qPCR of mouse GAPDH gene and *Ct* 16S rRNA gene. Ratio of 16S rRNA/mouse GAPDH was used to assess bacterial burden.

### ROS determination

Infected/control cells were grown in colourless media. 6 h after infection, Cell Rox deep red reagent was added according to manufacture’s protocol. ROS levels were detected using SpectraMax multi-mode plate reader, Molecular devices, USA at 644/655nm.

### NO levels determination

Infected/control cells were grown in colourless media for 6 h and media was collected to determine NO levels. NO levels were determined by mixing equal amounts of Griess reagent (Sigma-Aldrich) with supernatant and incubated for 10 min before taking O.D at 540nm. O.Ds were compared with 70 µg/ml of standard Nitrate.

### Arginase activity

Arginase activity was measured protein extract as mentioned earlier [34]. Enzyme activity was analysed by measuring the conversion of arginine to urea. Protein extract was incubated in 0.5 M L-arginine at 56°C for 10 min in presence of 2mM MnCl_2_ (pH 8). Reaction was stopped with stop solution (H_2_SO_4_/H_3_PO_4_/H_2_0 (1/3/7, v/v/v)), urea formation was measured by adding 40 ul α-isonitrosopropiophenone and heating at 95°C for 30 min. O.D was measured and enzyme activity was plotted as formation of 1 umol of urea per min per ug of protein.

### ELISA

Supernatants of macrophages were collected at 24hrs or otherwise mentioned time points after treatment. For T cell cytokines, APC-T cell co-culture supernatants were collected after 48hrs. Serum was separated from blood collected from infected/control animals and used to analyse the serum cytokine/Immunoglobulin levels. Vaginal lavage (VL) collected from mice of control and infected groups were also used to analyse cytokines. All the ELISAs were performed according to manufacture’s protocols (Supplementary table 2).

### Serum Ct-specific IgG assessment

10^4^ McCoy cells were plated in 96 well plate. Over-night grown cells were infected with *Ct* and after 24 h, cells were fixed with 1.5% PFA for 15 min. Cells were washed with PBS and blocked with 2.5% BSA in PBS for 1 hr. Serum separated from blood of mice from different experimental groups were added in 1:25 and 1:100 dilutions to plates and incubated for over-night at 4°C. Excess antibody was washed with PBS containing 0.5% Tween-20. Biotin-labelled anti-mouse IgG was added to wells and incubated for 1 h. After washing, streptavidin-HRP was added and incubated for 30 min. Excess strep-HRP was washed and bound antibody was measured using TMB conversion to coloured product. Readings were measured at OD450nm and plotted as relative IgG levels.

### Western Blot

Cell lysates were prepared in RIPA buffer containing protease inhibitor cocktail (Roche) and PMSF. 25ug of lysate was subjected to SDS-PAGE and transferred onto PVDF membrane (Millipore). Specific proteins were detected using Abs as listed in supplementary table 2. Protein loading was normalised using a mAb against GAPDH. Appropriate HRP-labelled secondary antibodies against Mouse IgG, Rabbit IgG and Sheep IgG were used to detect primary antibodies. ECL substrate was used to detect the secondary antibodies. Images were captured in FUSION-SL gel documentation system (Vilber, France).

### Quantitative real time (RT)-PCR

To compare gene expression in various samples, total RNA was extracted using TRIzol reagent and retrotranscribed using cDNA synthesis kit. Oligonucleotides for selected genes were designed and used for quantitative real-time PCR (given in supplementary table 3). Assays were performed in triplicates and results normalized according to the expression levels of GAPDH mRNA. Results were expressed using ΔΔC_T_ method for quantification.

### Flowcytometric analysis

Single cell suspensions from lymph nodes or activated T cells were prepared as described above. Flowcytometric analysis was done as described previously [36]. Cells were labelled with the following fluorescently labelled mAb as mentioned in supplementary table 2. For samples requiring intra-cellular labelling, cells were treated with brefeldin A after 6 hrs of stimulus and washed with wash buffer (BD Biosciences). Flowcytometric acquisition was done either in Attune NXT flowcytometry (ThermoFisher, USA) or CytoFlex (Beckman Coulter, USA) and analysed using Flowjo software.

### B cell proliferation analysis

Splenocytes were purified from naïve mice as mentioned earlier. RBC lysed splenocytes were passed through nylon wool column. After depletion of T cells, fresh media was added to column and agitated, cells were collected for B cells. Proliferation was measured by MTT assay after experimentation.

### Statistical tests

One-way ANOVA was used to analyse the significance between the groups. Two-way ANOVA was used to analyse the difference among subgroups and the difference among groups. Significance was denoted as p. (*** means p<000.1; ** means p<0.005; 8 means p<0.01). In figure 1, 2 and 4 # was used instead of * to denote the significance difference between groups.

## Data Availability

All data generated or analysed during this study are included in this published article and its corresponding supplementary file.

## Acknowledgment

Authors thank Dr. Bhaskar Saha, NCCS, Pune for critical evaluation of the manuscript. Acknowledgment to Dr. Manish Nivsarkar and Mr. Nikunj Tandel for helping in some parts of the animal experiments. Authors acknowledge Shivani Yadav, Parameswar Dalai, Hima Vora, Pranitha Rokkam, Devangi Zala, and Rakesh for their support in routine lab work. Part of the results presented in this are article are presented at 2020 Annual meeting of The American Society for Cell Biology.

## Author Contribution

NC performed the designed and performed the experiments and helped in drafting the manuscript. DS helped in performing the experiments. SKD extended animal facility. RAR conceived the study, designed the experiments and prepared and finalised the manuscript.

