## Supplementary figure for "IFNγ insufficiency during mouse intravaginal *Chlamydia trachomatis* infection exacerbates alternative activation in macrophages with compromised CD40 functions"

### Supplementary Figure (SF)

SF.1

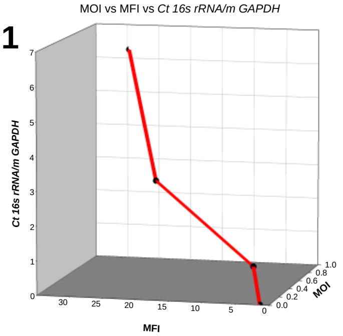


SF.2

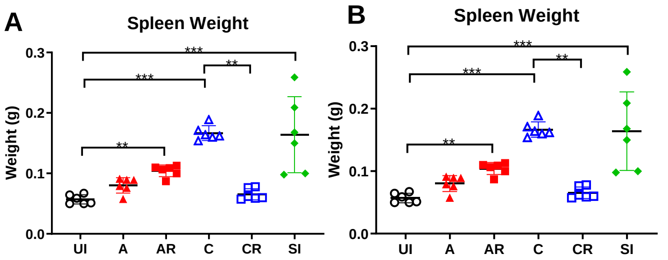


**Supplementary Figure 1:** Mouse fibroblast cells were infected with C. trachomatis at various doses and analysed for bacterial burden using Chlamydia OMP immunofluorescence and genomic content. MFI was calculated from confocal images and genomic content by qPCR analysis Ct 16s rRNA gene content from DNA. Graph was plotted to establish corelation

**Supplementary Figure 2:** Mice were infected with C.tr as indicated in Figure 1A and spleen weight (A) and inguinal lymph node weights (B) were measured after clearing of extra residual fat.

SF.3A

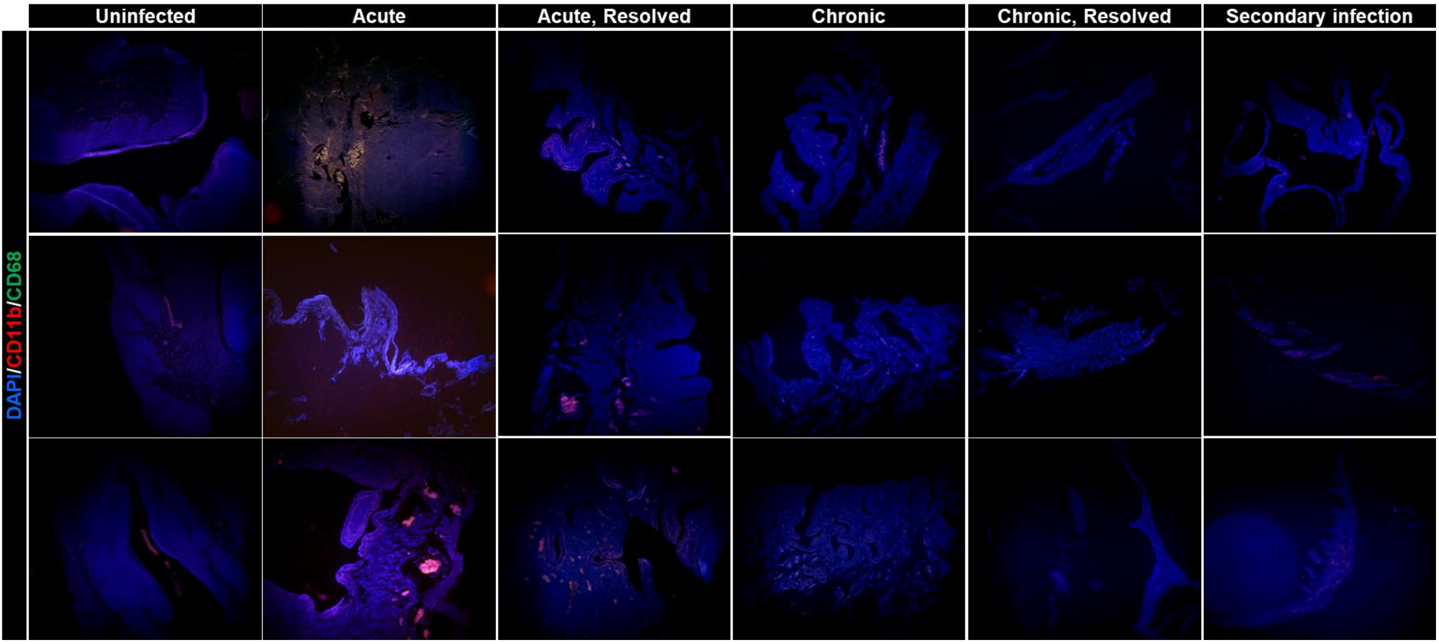


SF.3B

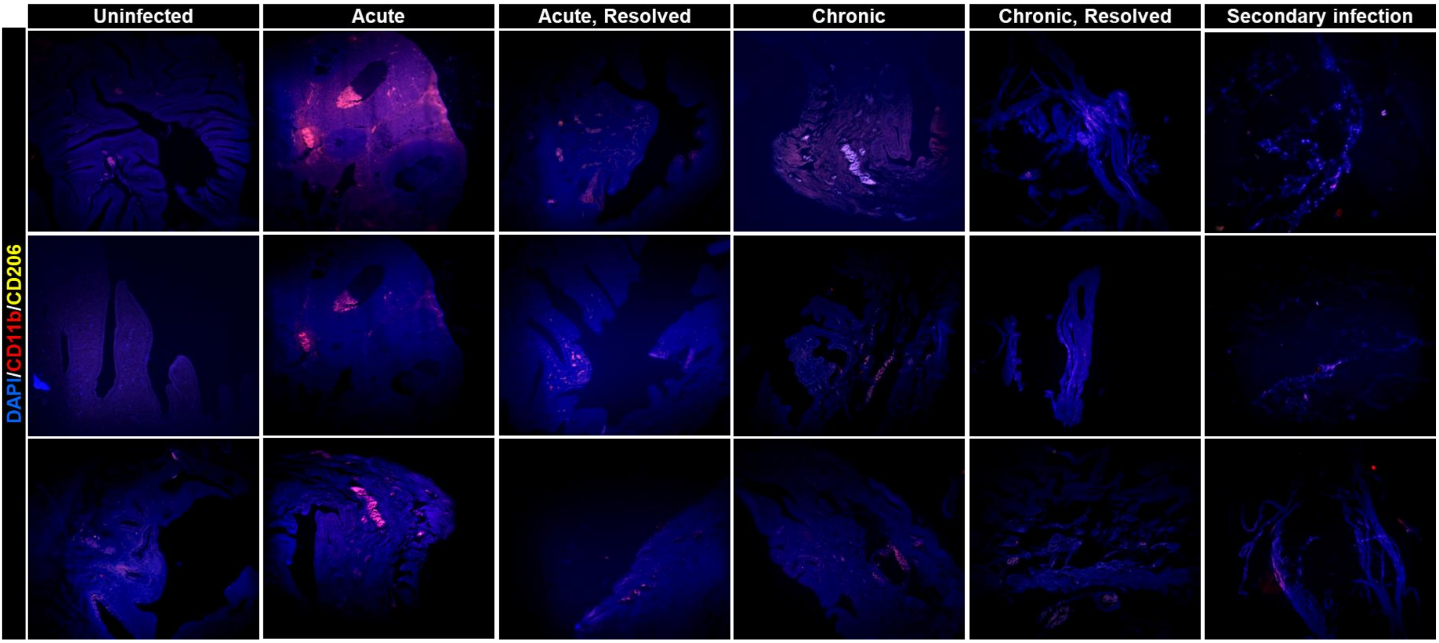

SF.3C

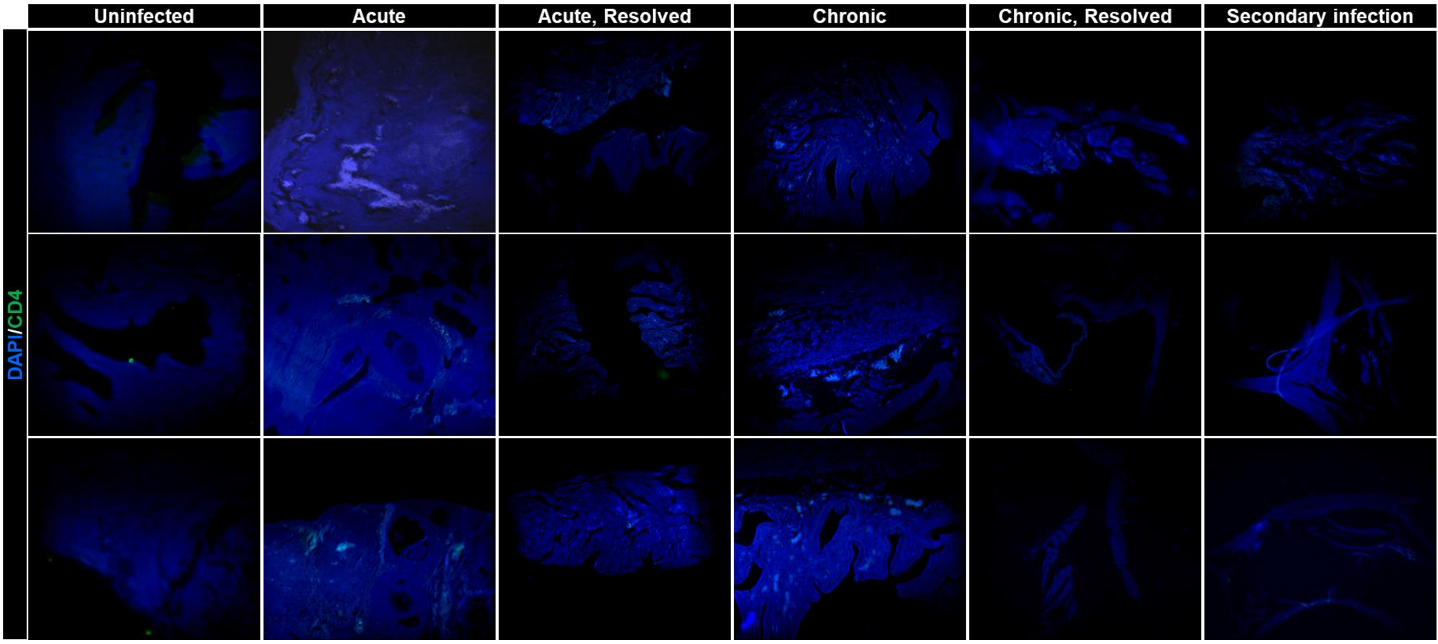


SF.3D

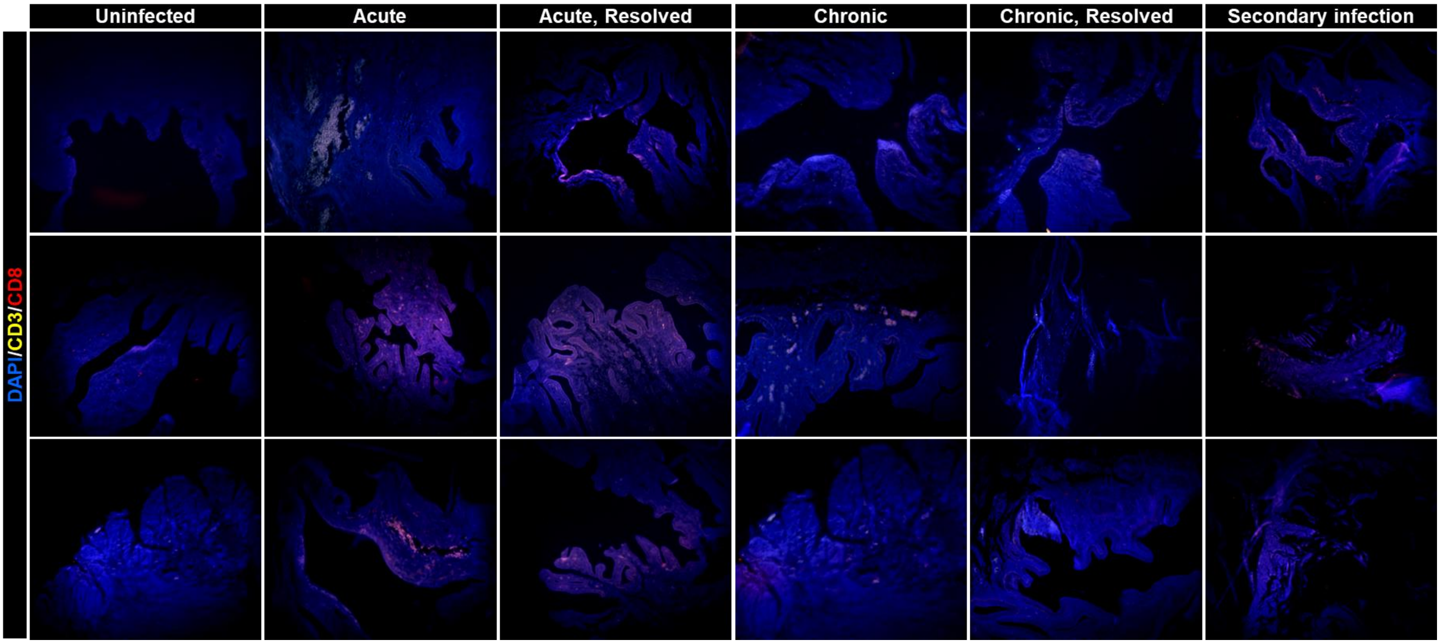


**Supplementary Figure 3:** (A) Tissue section were stained for CD68, CD11b and nuclear material with rabbit anti-CD68, anti-CD11b labelled with APC and DAPI. Anti-rabbit IgG Alexa 488 was used to detect anti-CD68 antibody. Sections were observed in GFP, RFP and DAPI filter in Olympus fluorescence microscope under margination of 200X. (B) Tissue section were stained for CD206, CD11b and nuclear material with sheep anti-CD68, anti-CD11b labelled with APC and DAPI. Anti-sheep IgG Alexa 568 was used to detect anti-CD68 antibody. Sections were observed in GFP, RFP and DAPI filter in Olympus fluorescence microscope under margination of 200X. (C) Tissue section were stained for CD4 in the background of nuclear stain DAPI and was detected using anti-rabbit IgG labeled with FITC. Sections were observed in GFP, RFP and DAPI filter in Olympus fluorescence microscope under margination of 200X. (D) Immunohistoflourescene was performed for CD3 and CD8 using rabbit anti-mouse CD3 and anti-mouse CD8 labelled with APC. Anti-CD3 was detected using anti-rabbit IgG labelled with Alexa 568. Merged images were presented in Figure 2G. Minimum of 6 unsupervised ROIs were captured for each section prepared from a mice for all the immunofluorescence experiments. All the images were processed blindly and merged in Image J, NIH USA.

SF.4

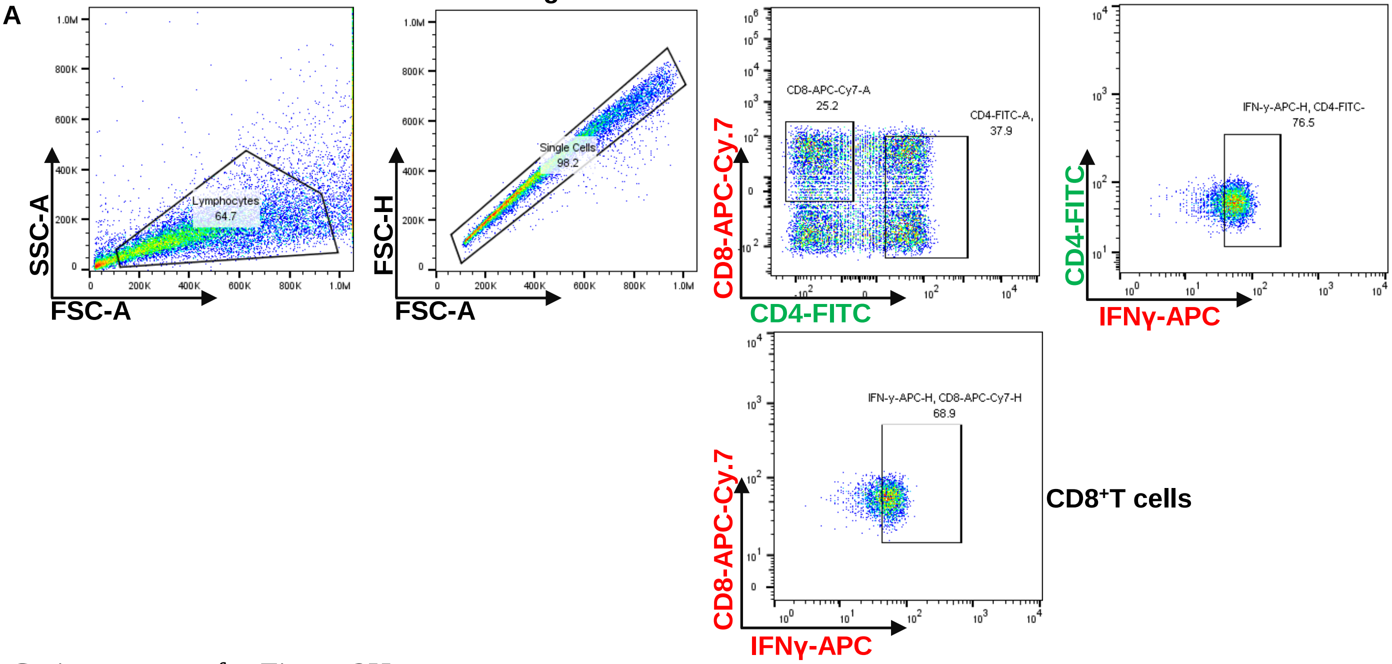


Gating strategy for Figure 2H.

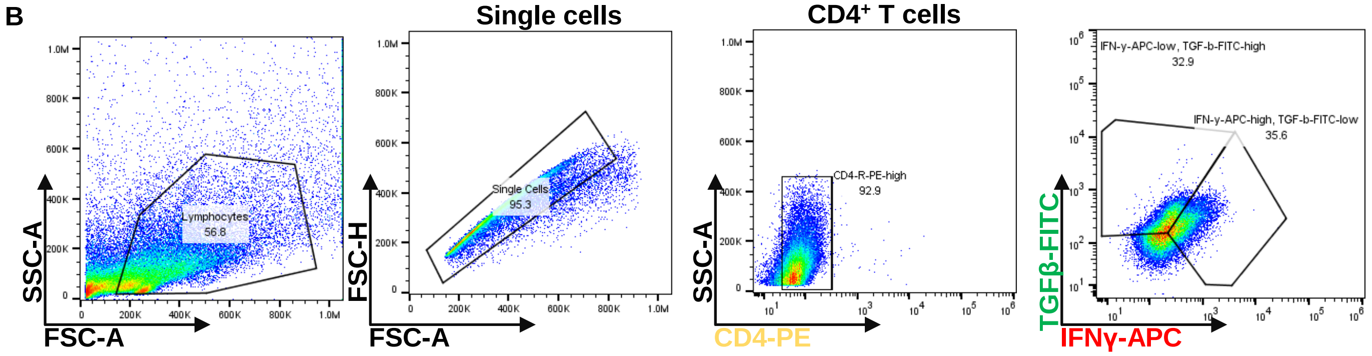


Gating strategy for Figure 2K

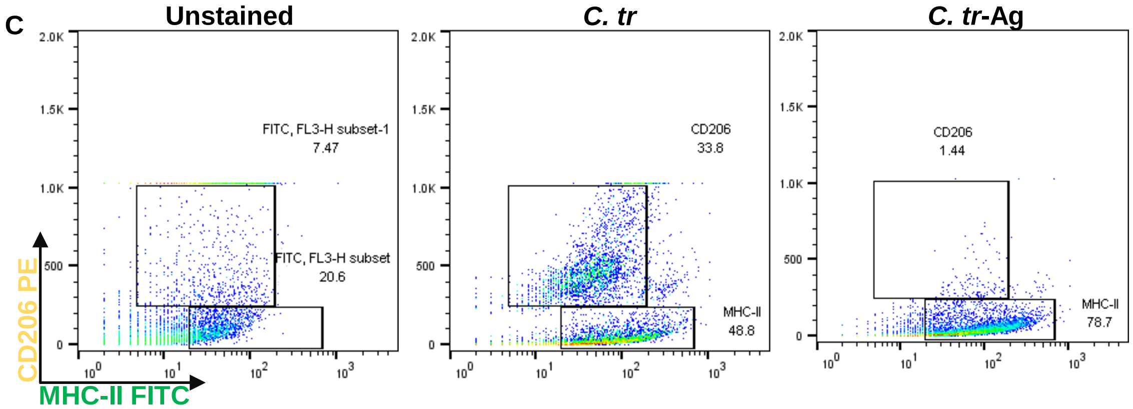


**Supplementary figure 4: Flowcytometric analysis.** (A) $CD3^+$ T cells isolated from lymph nodes were stained for CD4, CD8 and intracellular IFNγ and analyzed for CD4 and CD8 T cell distribution and $CD4^+IFN^+$- and $CD8^+$ $IFN^+$- T cells. % of cells are plotted in figure 2H. Scatter plot analysis of CD11b purified bone marrow macrophages stained for CD206 and MHC-II with anti-CD206 labelled with PE and anti-MHC-II labelled with FITC. % of cells were plotted in Figure 3H

SF5

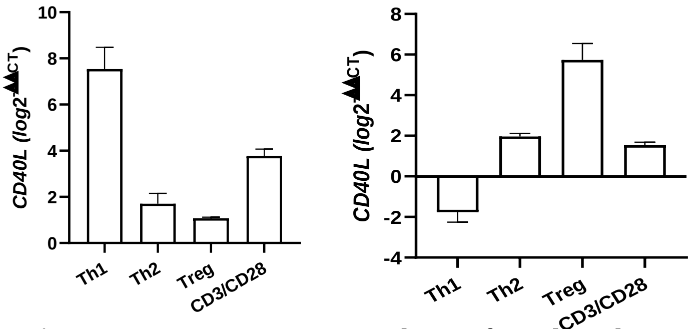


**Supplementary Figure 5:** BMDMs were either infected with *C.tr* or treated with *C.tr*-Ag for 24hrs and stained for CD206 and MHC-II with anti-CD206 labelled with PE and anti-MHC-II labelled with FITC

SF6

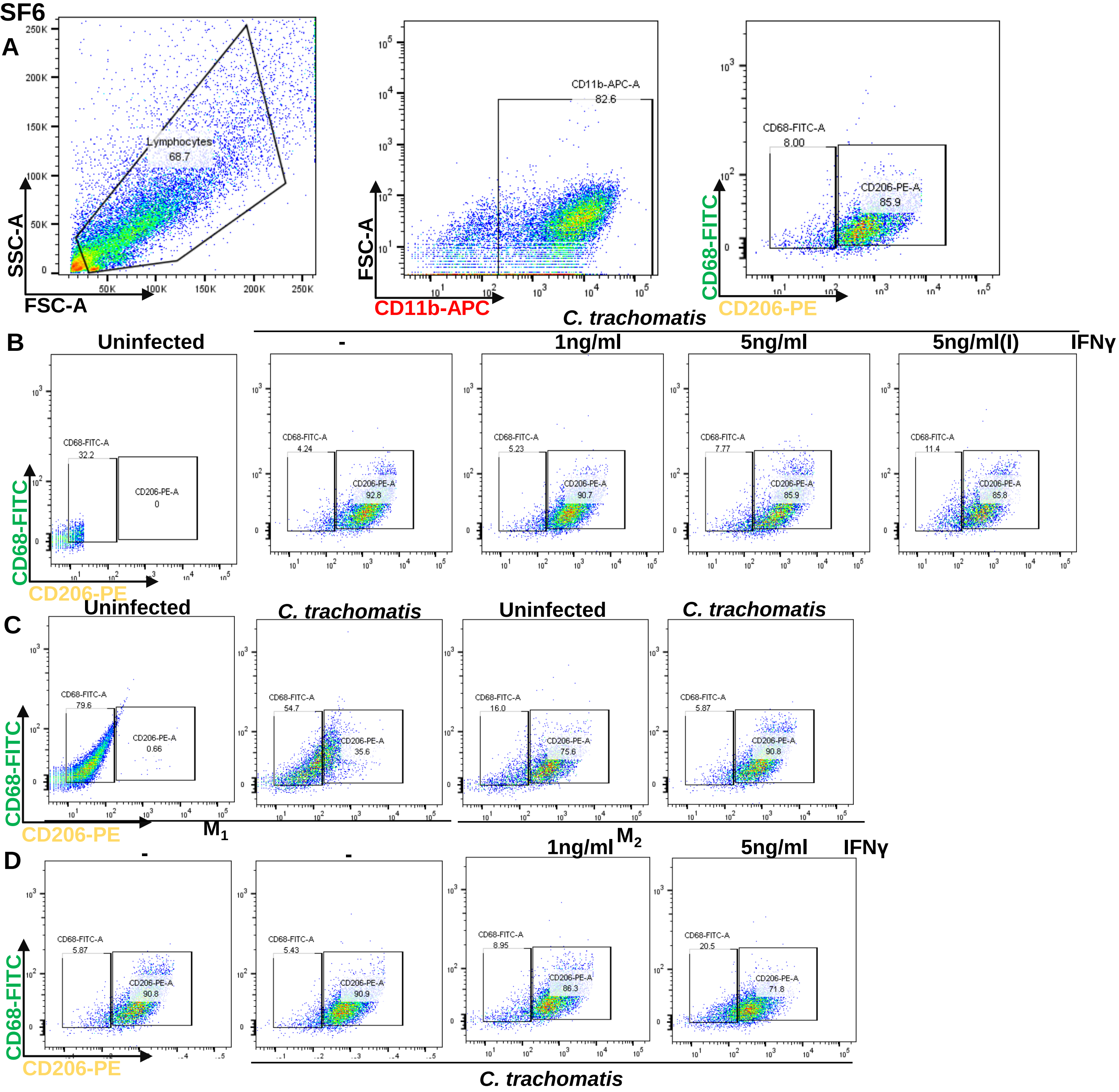


**Supplementary figure 6: Flowcytometric analysis.** (A) Gating strategy for macrophage polarization analysis i.e., CD206 and CD68 expression of CD11b$^{+}$ cells. (B) Scatter plot analysis of CD11b$^{+}$ bone marrow derived macrophages for CD206 and CD68 expression. % of cells were plotted in Figure 7F. (C)$M_1$(GM-CSF and IFNγ) and $M_2$(MCSF, IL-10 and IL-4) derived cells infected wit C.tr were analyzed for macrophage polarization. % of CD11b cells expressing CD206 and CD36 were plotted in Figure 7D. (D) $M_2$ were infected and supplemented with IFNγ (1ng/ml and 5ng/ml) and analysed for macrophage polarization (Figure 7G).

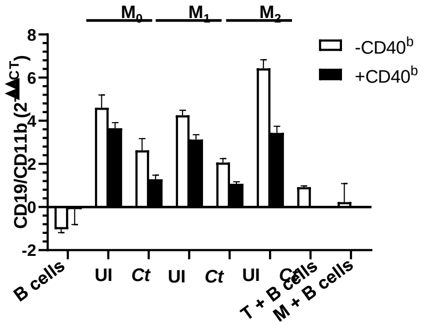


**Supplementary figure 7: B cell proliferation.** Total RNA from the experiment Fig.8L was extracted and analyzed for CD19 expression (B cell marker) and CD11b expression (macrophage marker) and the ratio of them was plotted to indicate the B cell proliferation.
