## Supplementary table for "IFNγ insufficiency during mouse intravaginal *Chlamydia trachomatis* infection exacerbates alternative activation in macrophages with compromised CD40 functions"

**Supplementary table 1:** Recombinant proteins

| **Recombinant protein** | **Concentration** | **Catalogue no.** | **Source** |
| --- | --- | --- | --- |
| **GM-MCSF** | 20 ng/ml | 554586 | BD Pharmigen, USA |
| **M-CSF** | 20 ng/ml | 14-8983-80 | eBioscienes, ThermoFisher Scientific, USA |
| **IL-4** | 10 ng/ml | 550067 | eBioscienes, ThermoFisher Scientific, USA |
| **IL-10** | 10 ng/ml | 550070 | BD Pharmigen, USA |
| **IFNγ** | 10 ng/ml | 554587 | BD Pharmigen, USA |
| **TGFβ** | 10 ng/ml | 100-21 | Peprotech, USA |
| **IL-2** | 10 ng/ml | 550069 | BD Pharmigen, USA |
| **EGF** | 10 ng/ml | 315-09 | Peprotech, USA |
| **Insulin** | 5 ug/ml | 11070-73-8 | Sigma-Aldrich, USA |

**Supplementary Table 2: Antibodies used in the study**

| **Unlabelled antibodies used in immunoblotting (1:1000) and IHC (1:250)** | | | | | | | | | | | | | |
| --- | --- | --- | --- | --- | --- | --- | --- | --- | --- | --- | --- | --- | --- |
| **Antigen** | | | | | **Isotype** | | | **Molecular weight (KD)** | | **Brand** | | **Catalogue No.** | |
| **GAPDH** | | | | | Mouse IgG | | | 37 | | Invitrogen, Thermo Fisher, USA | | AM | |
| **NOS II** | | | | | Rabbit IgG | | | 125 | | Merck Millipore, USA | | 10-3018 | |
| **ERK1/2(Pt185/Py187)** | | | | | Rabbit IgG | | | 44 | | Invitrogen, Thermo Fisher, USA | | 700031 | |
| **Total ERK1/2** | | | | | Rabbit IgG | | | 44/42 | | Invitrogen, Thermo Fisher, USA | | AHR0024 | |
| **Phospho p38 β2** | | | | | Mouse IgG2 | | | 38 | | Invitrogen, Thermo Fisher, USA | | 700012 | |
| **P38 MAPK** | | | | | Rabbit IgG | | | 38 | | Invitrogen, Thermo Fisher, USA | | AHO1362 | |
| **CD40** | | | | | Rabbit IgG | | | 43 | | Invitrogen, Thermo Fisher, USA | | PA5-78980 | |
| **CD68** | | | | | Rabbit IgG | | |  | | Sigma-Aldrich | | SAB5700832 | |
| **CD206** | | | | | Goat IgG | | |  | | R & D Systems, Biotechne, USA | | AF5868 | |
| **Flowcytometry antibodies** | | | | | | | | | | | | | |
| **Antigen** | | **Conjugate** | **Isotype** | | | | **Brand** | | | | | | **Catalogue No.** |
| **CD16/32** | | Fc block |  | | | | BD Pharmingen, USA | | | | | | 553141 |
| **CD11b** | | APC | Rat IgG2 | | | | BioLegend, USA | | | | | | 101211 |
| **CD68** | | PE | Rat IgG2a | | | | BD Pharmigen, USA | | | | | |  |
| **CD206** | | Alexa 488 | Rat IgG2 | | | | eBioscience, Thermo Fisher, USA | | | | | | 53-2061-80 |
| **CD4** | | FITC | Rat IgG2b | | | | eBioscience, Thermo Fisher, USA | | | | | | 11-0042-81A |
| **CD8** | | APC-Cy7 | Rat IgG2a | | | | BD Pharmigen, USA | | | | | | 561967 |
| **IFNγ** | | APC | Rat IgG1 | | | | eBioscience, Thermo Fisher, USA | | | | | | 17-7311-81 |
| **IL-10** | | Alexa 488 | Rat IgG2b | | | | eBioscience, Thermo Fisher, USA | | | | | | 53-7101-80 |
| **IL-4** | | PE | Rat IgG1 | | | | Biogems, Peprotech, USA | | | | | | 49-9856 |
| **Foxp3** | | PE | Rat IgG2b | | | | eBioscience, Thermo Fisher, USA | | | | | | 12-5773-80A |
| Isotype controls | | | | | | | | | | | | | |
| **Rat IgG1** | | APC |  | | | | BD Pharmingen, USA | | | | | | 554686 |
| **Rat IgG1** | | PE |  | | | | BD Pharmingen, USA | | | | | | 551979 |
| **Rat IgG2** | | APC |  | | | | eBioscience, Thermo Fisher, USA | | | | | | 12-4321-81A |
| **Functional Antibodies** | | | | | | | | | | | | | |
| **Antigen** | Activiity | | | Isotype | | Clone | | | Brand | | | | Catalogue No. |
| **CD40^a^** | Stimulation | | | Rat IgG2a | | 3/23 | | | BD Pharmingen, USA | | | |  |
| **CD40^b^** | Blocking | | | Armenian Hamster IgM | | HM40-3 | | | BD Pharmingen, USA | | | | 553057 |
| **CD3** | Stimulation | | | Armenian Hamster IgG1 | | 145-2C11 | | | BD Pharmingen, USA | | | | 559532 |
| **CD28** | Stimulation | | | Syrian Hamster IgG | | 37.51 | | | BD Pharmingen, USA | | | | 557401 |
| **Secondary Antibodies** | | | | | | | | | | | | | |
| **Anti-Rabbit-IgG** | | | | Biotin | | | | Novex, Thermo Fisher, USA | | | | | AHO1202 |
| **Ant-Rabbit IgG** | | | | HRP | | | | ThermoFisher Scientific, USA | | | | | 656120 |
| **Anti-Rabbit IgG** | | | | Alexa 488 | | | | ThermoFisher Scientific, USA | | | | | A11034 |
| **Anti-Rabbit IgG** | | | | Alexa 546 | | | | ThermoFisher Scientific, USA | | | | | A11010 |
| **Anti-Rat-IgG** | | | | Alexa 568 | | | | ThermoFisher Scientific, USA | | | | | A11077 |
| **Anti-Mouse IgG** | | | | HRP | | | | ThermoFisher Scientific, USA | | | | | A16072 |
| **Anti-Mouse IgG** | | | | Alexa 488 | | | | ThermoFisher Scientific, USA | | | | | A11029 |
| **Anti-Goat IgG** | | | | Biotin | | | | ThermoFisher Scientific, USA | | | | | PA1-28663 |
| **Streptavidin** | | | | PE | | | | BioLegend, USA | | | | | 405203 |
| ELISA Paired antibodies (Coating 1:250; Detection- 1:500) | | | | | | | | | | | | | |
| **IL-1β** | Coating | | | BD OptEIA | | | | | | | 559603 | | |
|  | Biotin | | |  |  |  |  |  |  |  |  |  |  |
| **IL-12** | Coating | | | eBiosciences, ThermoFisher Scientific, USA | | | | | | | 14-7125-81 | | |
|  | Biotin | | | eBiosciences, ThermoFisher Scientific, USA | | | | | | | 13-7123-81 | | |
| **IL-10** | Coating | | | eBiosciences, ThermoFisher Scientific, USA | | | | | | | 14-7102-67B | | |
|  | Biotin | | | eBiosciences, ThermoFisher Scientific, USA | | | | | | | 554423 | | |
| **TGF-β** | Coating | | | BD Biosciences, USA | | | | | | | 555052 | | |
|  | Biotin | | | BD Biosciences, USA | | | | | | | 555053 | | |
| **TNF-α** | Coating | | | BD Biosciences, USA | | | | | | | 3207537 | | |
|  | Biotin | | | BD Biosciences, USA | | | | | | | 3331772 | | |
| **IL-4** | Coating | | | eBiosciences, ThermoFisher Scientific, USA | | | | | | | 14-7041-67B | | |
|  | Biotin | | |  |  |  |  |  |  |  | 13-7042-67D | | |
| **IL-2** | Coating | | | eBiosciences, ThermoFisher Scientific, USA | | | | | | | 14-7022-67B | | |
|  | Biotin | | | eBiosciences, ThermoFisher Scientific, USA | | | | | | | 13-7021-67D | | |
| **IL-17** | Coating | | | eBiosciences, ThermoFisher Scientific, USA | | | | | | | 560268 | | |
|  | Biotin | | | eBiosciences, ThermoFisher Scientific, USA | | | | | | | 555067 | | |
| **IFN-γ** | Coating | | | BD OptEIA | | | | | | | 555138 | | |
|  | Biotin | | |  |  |  |  |  |  |  |  |  |  |
| **Avidin-HRP** | | | | BD Pharmigen, USA | | | | | | | 554066 | | |

**Supplementary table 3: Primer sequences**

| Gene |  | Sequence (3′-5′) |
| --- | --- | --- |
| *PD1* | Forward | CCAGTCAGTGGAGAGTGGAG |
|  | Reverse | GCAGCATTCGTAGAATAACCCT |
| *Tim3* | Forward | GGGAGGACAGTCGTCACCAA |
|  | Reverse | GGGAGCCAGCACAGATCAAG |
| *KLRG1* | Forward | CCTTACATTTCCGGACAACCA |
|  | Reverse | CCCACCTCCAGCCATCAAT |
| *Lag3* | Forward | GCCCTGTGCTGGAGATTCA |
|  | Reverse | GCTAGACTCTGCGGCGTACAC |
| *CD57* | Forward | AGGCTGCCTTCTTAGCCAAAC |
|  | Reverse | CTGCCAAAACGTGAAAAGTAAAAA |
| *CTLA4* | Forward | GTGAACCTCACCATCCAAGGA |
|  | Reverse | TGCCCATGCCCACAAAGTAT |
| *Arginase-1* | Forward | CTCGAGGTCCTGAGTTGCAC |
|  | Reverse | GCTTCTTCTGTCCCCGAGAG |
| *IFNγR I* | Forward | TACAGGTAAAGGTGTATTCGGGT |
|  | Reverse | ACCGTGCATAGTCAGATTCTTTT |
| *IFNγR II* | Forward | CTTCTCCCCTCCCTTTGATGT |
|  | Reverse | AGGGCCTTCAACCTGTTCCT |
| *IDO* | Forward | GGCTTCTTCCTCGTCTCTCTATTG |
|  | Reverse | CTTTCAGGTCTTGACGCTCTACTG |
| *MHC-I^h2k^* | Forward | CAGAGAGCCAAGAGCGATGAG |
|  | Reverse | TCCGCTGGAACGTGTGAGA |
| *Erg2* | Forward | CAGTTCAACCCCTCTCCAAAAA |
|  | Reverse | CTTGGCGGTCATCATTTGCT |
| *C/EBPβ* | Forward | CTGCCAAAACGTGAAAAGTAAAAA |
|  | Reverse | AACCCCGCAGGAACATCTTT |
| *CD40L* | Forward | TGGTAATGCTTGAAAATGGGAAA |
|  | Reverse | CTCGAAGGCTCCCGATTAGA |
| *Granzyme B* | Forward | CCCAGGCGCAATGTCAAT |
|  | Reverse | CCCCAACCAGCCACATAGC |
| *Perforin* | Forward | AGAGGAAGTTCTGCGTTTTACCA |
|  | Reverse | CTGTCGGATGGGACAACCA |
| *CD206* | Forward | GGCTGATTACGAGCAGTGGA |
|  | Reverse | ATGCCAGGGTCACCTTTCAG |
| *CD36* | Forward | ATGGGCTGTGATCGGAACTG |
|  | Reverse | GTCTTCCCAATAAGCATGTCTCC |
| *iNOS* | Forward | GGATCTTCCCAGGCAACCA |
|  | Reverse | CAATCCACAACTCGCTCCAA |
| *T-bet* | Forward | ACCTGTTGTGGTCCAAGTTCAA |
|  | Reverse | GCCGTCCTTGCTTAGTGATGA |
| *PDI* | Forward | CAGCCAATGATGTGCCTTCTC |
|  | Reverse | TTTAATTCACGGCCACCTTCA |
| *Ehrin B2* | Forward | CTTCCCCCTCCCTGATTGA |
|  | Reverse | TCTAAGAGGTGGGCCATGTGT |
| Primers for genomic DNA amplification | | |
| *Mouse GAPDH* | Forward | CCGCCATGTTGCAAACG |
|  | Reverse | CGAGAGGAATGAGGTTAGTCACAA |
| *C.tr 18S rRNA* | Forward | ATGCCGCCTGAGGAGTACAC |
|  | Reverse | TTCGCGTTGCATCGAATTAA |
